# Cellular context restricts a promiscuous m6A reader IDR to a single functional effector interface for mRNA decay

**DOI:** 10.64898/2026.08.04.742832

**Authors:** Wei Yee Chan, Enkai Jin, Hector Cross, Laura Masino, Fairouz Ibrahim, Waleed S Albihlal, Duncan Berger, James Boot, Aleksandra Michrowska, Stephane Mouilleron, Mark Skehel, Kamil R Kranc, Folkert J van Werven

## Abstract

*N6*-methyladenosine (m6A) is a conserved mRNA modification that regulates transcript stability, yet how m6A readers engage effectors remains unknown. We show that during yeast meiosis, the YTH-domain protein Pho92 promotes the decay of m6A-modified transcripts through an uncharacterised, promiscuous intrinsically disordered region (IDR). *In vitro*, the Pho92 IDR makes multiple contacts with the Ccr4-NOT deadenylase complex and drives tethered reporter decay in the absence of any single subunit. Yet in meiosis, turnover of endogenous m6A-modified transcripts strictly depends on a single interface: direct binding of the Pho92 IDR to Caf40. A single hydrophobic residue substitution is sufficient to abolish Caf40 binding and halt transcript decay. Human YTHDF proteins can functionally substitute for Pho92, but require multiple hydrophobic patches within the IDR for decay activity. Thus, on endogenous m6A-modified transcripts during meiosis, the promiscuous Pho92 IDR is restricted to a single functional interface, revealing unappreciated specificity in m6A-directed mRNA decay.

## Introduction

Post-transcriptional regulation is a fundamental layer of gene expression control, enabling cells to rapidly adjust mRNA stability and translation in response to developmental and environmental cues. Among the mechanisms that regulate mRNA fate, chemical modifications of mRNA have emerged as key determinants of post-transcriptional gene expression. *N6*-methyladenosine (m6A) is the most abundant internal modification on eukaryotic mRNA, and its deposition by the m6A methyltransferase complex (MTC) at consensus motifs within 3’ UTRs and open reading frames regulates diverse aspects of mRNA metabolism, including stability, translation, and splicing^1^.

The biological consequences of m6A are mediated by YTH domain-containing reader proteins, which selectively recognize m6A-modified transcripts via their conserved YTH domains. In mammals, the cytoplasmic readers YTHDF1, YTHDF2, and YTHDF3 have been shown to control the stability and translation of m6A-modified transcripts^2,3^. Among these, m6A-enhanced mRNA decay is perhaps the best-characterized outcome, with m6A-modified transcripts exhibiting substantially higher turnover rates than their unmodified counterparts. Mechanistically, YTHDF proteins are thought to recruit effector complexes that promote mRNA degradation, including the Ccr4-NOT deadenylase complex and components of the nonsense-mediated decay pathway^4,5^. However, the molecular interfaces that govern effector recruitment remain poorly defined, in part because YTHDF paralogs are functionally redundant in mammalian cells, making it difficult to discriminate these processes^2^. The budding yeast *Saccharomyces cerevisiae* provides a powerful model to address this problem^6^. In yeast, the m6A pathway is active specifically during meiosis, where the MTC deposits m6A on more than 1000 mRNAs as part of the sporulation program^7,8^. Yeast encodes a single YTH domain reader, Pho92 (also known as Methylated RNA-binding protein 1), which is essential for the timely progression through meiosis^7,9,10^. Pho92 promotes the turnover of m6A-modified transcripts, using a mechanism that requires translation^9,10^. This is consistent with emerging models in human cells linking m6A within open reading frames to ribosome-associated mRNA decay^11,12^. The m6A pathway in yeast, with a single reader, a defined set of targets, and a restricted developmental window, offers an opportunity to dissect the molecular regulatory mechanisms of m6A RNA biology.

Here, we examined the mechanism by which the YTH domain reader, Pho92, promotes the turnover of m6A-modified transcripts. Through proximity labelling, structural modelling, and functional analysis, we show that Pho92 interacts with the Caf40 subunit of the Ccr4-NOT complex via its uncharacterized N-terminal intrinsically disordered region (IDR). The Pho92 IDR can contact multiple interfaces of Ccr4-NOT *in vitro* and in a heterologous reporter. However, a single hydrophobic residue within the IDR is essential for Caf40 interaction and the turnover of m6A-modified transcripts in meiosis. Human YTHDF proteins can functionally substitute for Pho92 and similarly require hydrophobic IDR residues for decay activity, indicating partial mechanistic conservation. Thus, despite its promiscuous binding capacity, the Pho92 IDR is constrained to a single functional interface for the decay of endogenous m6A-modified transcripts during yeast meiosis.

## Results

### Proximity of Pho92 identified candidate effectors

To gain insight into the Pho92 interactome in cells, we developed an improved proximity labelling protocol. In short, we tagged Pho92 with the TurboID enzyme (Pho92-TID) at the endogenous locus^13,14^. We induced cells to enter meiosis. Briefly, we first grew cells to saturation in rich medium, overnight in pre-sporulation medium, and subsequently shifted cells into sporulation medium for 4 hours in the presence of biotin. We then fixed cells with trichloroacetic acid (TCA) and prepared denaturing protein extracts, from which we pulled down biotinylated proteins using streptavidin-coated beads (see Materials and Methods for details) (Figure S1A and S1B). The eluates were analysed by mass spectrometry.

Using the optimized proximity labelling protocol, we validated that the TID enzyme (on Pho92-TID) biotinylated Pho92 by western blot (Figure S1A). In addition to the endogenously biotinylated proteins (background), we detected unknown biotinylated proteins in the Pho92-TID eluate, indicating that Pho92-TID biotinylated proximal interactors (Figure S1C and S1D). Notably, the TID did not affect Pho92 function, because Pho92-TID cells entered meiosis with similar kinetics as the wild type (Figure S1E). To assess whether the labelling was dependent on m6A, we also performed proximity labelling with Pho92-TID in the *slz1*Δ mutant. Slz1 is a subunit of MTC required for m6A deposition. Pho92 cannot stably associate with mRNAs in the *slz1*Δ mutant, allowing us to determine whether interactions are RNA-dependent (Figure 1A).

**Figure 1.**
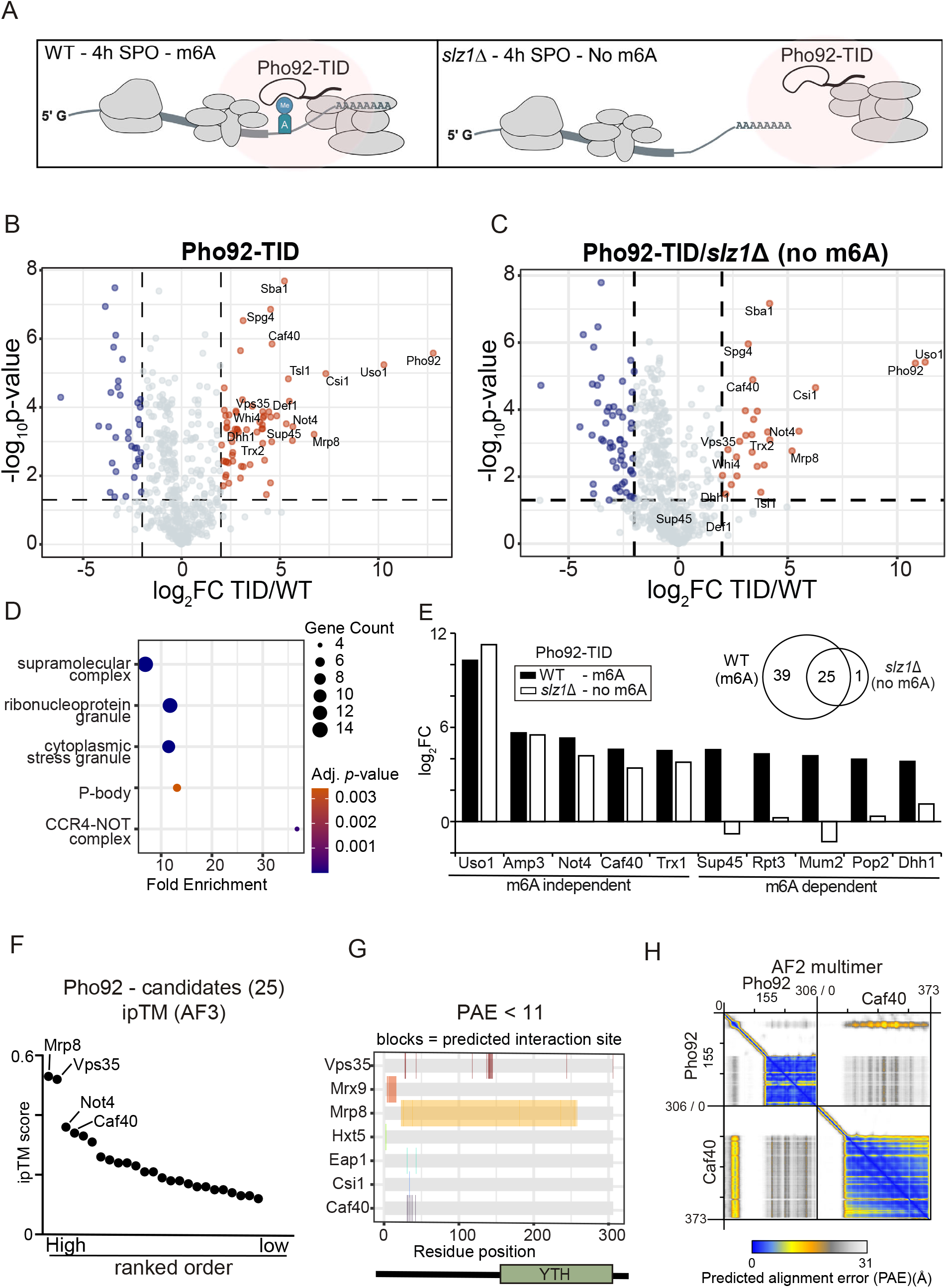
m6A-dependent and m6A-independent proximity labelling of Pho92. **(A**) Schematic cell setup of the proximity labelling using the TurboID experiment of Pho92 (Pho92-TID) in wild-type and *slz1*Δ cells staged in early meiosis. Cells were induced to enter meiosis by growing till saturation in rich medium (YPD), overnight in pre-sporulation medium, and subsequently for 4 hours in sporulation medium (SPO) plus biotin. Cells were fixed with TCA, and proteins were solubilised with SDS sample buffer. Biotinylated proteins were enriched with streptavidin beads, and LC-MS analysed eluates. The *slz1*Δ has no detectable m6A and serves to identify m6A-independent protein interactions. (**B**) Volcano plot of Pho92-TID (FW11105) versus wild-type control (FW1511). The x-axis represents fold change (log2 FC), and the y-axis -log t-test p-value. Significantly enriched proteins are colored in red. (**C**) Volcano plot comparing Pho92-TID profiles in *slz1*Δ background (FW11200) to wild-type control. (**D**) Gene ontology (GO) analysis of enriched proteins of the proximity labelling experiments of B. (**E**) Selection of enriched proteins in the Pho92-TID that show m6A-dependent or independent enrichment. Also shown is a Venn diagram of the overlap between m6A-dependent or independent enriched proteins. (**F**) Ranked analysis of AlphaFold 3 predictions between 26 m6A-independent enriched proteins in TurboID from Figure 1E. Highlighted are predicted interactions with an ipTM value of 0.3 or higher. (**G**) Graphical outline of PAE analysis (see materials and methods) of proteins that showed an m6A-independent enrichment with Pho92-TID. We used PAE <11 as the cut-off. (**H**) AlphaFold2 multimer PAE plot of Pho92 and Caf40.

We found that 64 proteins were significantly enriched in Pho92-TID, and 25 were detected in *slz1*Δ cells (FC > 2, -log10 p < 1.3) (Figure 1B and 1C). The Pho92-TID detected proteins showed enrichment for components of stress granules (e.g. Dhh1), P-body (e.g. Tif4631), and more specifically, the Ccr4-NOT complex (e.g. Caf40, Not4, Not3) (Figure 1D). Of the 64 enriched proteins in the wild type, 39 were not enriched in *slz1*Δ cells, suggesting they were likely RNA-dependent interactions (Figure 1E). For example, the Mum2 subunit of the yeast MTC and translation termination factor Sup45 showed m6A-dependent labelling, suggesting that they co-occupy m6A-modified mRNAs with Pho92 (Figure 1E)^15^. The Caf40 and Not4 subunits of the Ccr4-NOT complex remained enriched in *slz1*Δ cells, indicating that these subunits are potentially newly identified direct interactors with Pho92 (Figure 1E and S1F).

Next, we used AlphaFold to examine whether enriched proteins with Pho92-TID that were not dependent on m6A modification formed a potential direct interaction with Pho92. We performed pairwise interaction analyses using AlphaFold Multimer and AlphaFold3^16,17^. We found that out of 25 potential interactions, 5 proteins had an ipTM value of 0.3 or higher, and 7 proteins had predicted interaction sites with predicted alignment error (PAE) of less than 11 and a distance <3Å (Figure 1F and 1G). While these cut-offs are relatively lenient, taking these two screening methods, only three proteins made both thresholds (Caf40, Mrp8 and Vps35). Mrp8 is a protein of unknown function and has been implicated in localising to mitochondria, while Vps35 is implicated in endosomal trafficking^18^. Thus, both are less likely to be involved in RNA decay. In contrast, Caf40 is a strong candidate regulator of the decay of m6A-modified transcripts because it is a key subunit of the Ccr4-NOT complex and functions as a scaffold for RNA-binding proteins that promote mRNA degradation^19–21^. Additionally, Caf40 and Pho92 yielded consistent structural interaction predictions in AlphaFold 2 multimer and AlphaFold 3 models (see methods and materials, Figure 1H and S1G). These analyses suggest that Caf40 is a potential direct interactor with Pho92.

### *In vitro*, Pho92 directly interacts with Caf40 via multiple sites in the N-terminal IDR

The predicted interaction between Caf40 and Pho92 is mediated via the convex surface of Caf40 and an N-terminal segment of Pho92 (Figure 2A). Structurally, Pho92 features a conserved C-terminal YTH domain that directly binds to m6A-modified transcripts. In contrast, its extensive N-terminal region (spanning approximately half of the total sequence) lacks globular domains. It is predicted to be largely intrinsically disordered (IUPred2) and has low pLDDT scores (AlphaFold 2) (Figures 2B and S2A)^22,23^. However, closer inspection of residues 28 to 52 within the IDR shows a marked dip in disorder score consistent with a transient or stable α-helical conformation (Figures 2A and 2B). AlphaFold2-Multimer predicts that this helical segment docks onto the concave surface of Caf40 (Figure 2A and S2A, S2B). The N-terminal IDR of Pho92 also contains two tryptophans (W5 and W18), of which W5 is predicted to interact in the hydrophobic pocket on the convex side of Caf40 (Figure 2A and Figure S2B). Thus, structural models suggest that an α-helical region and tryptophan residues in the Pho92 N-terminal IDR potentially mediate the interaction with Caf40.

**Figure 2.**
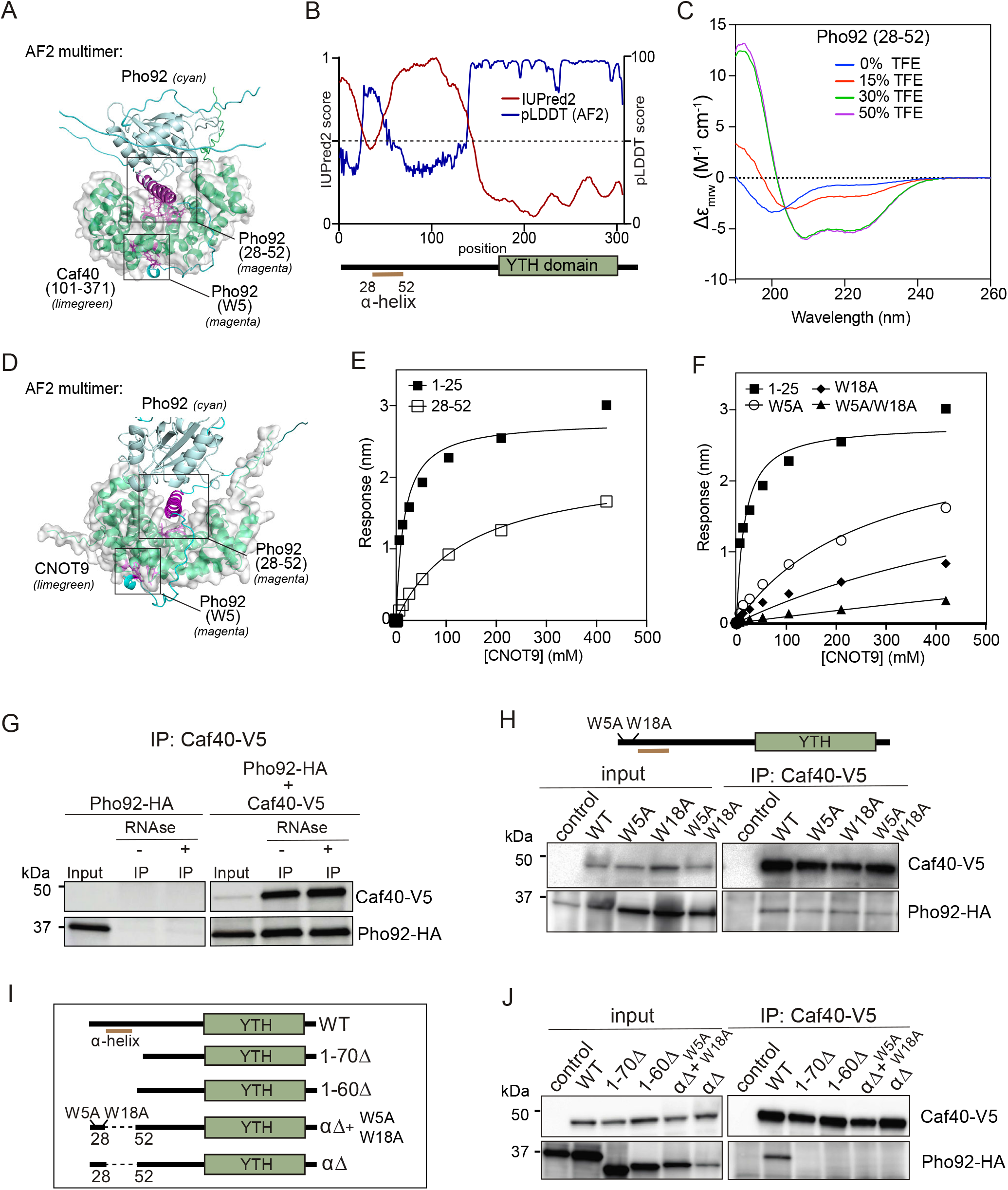
Pho92 and Caf40/CNOT9 interactions have distinct requirements *in vitro* and in cells. **(A)** Representative AlphaFold2 model highlighting Pho92 (cyan) and Caf40 (101-371, limegreen) with predicted interaction surfaces – Pho92 W5 (magenta) and Pho92 28-52 (magenta). See figure S2A and S2B for further details on predicted interaction surfaces and residues. (**B**) Pho92 intrinsically disordered region prediction using PrDOS (red). pLDDT score of Pho92 from Alphafold2 (blue). (**C**) Circular Dichroism (CD) spectroscopy of synthetic peptide (Pho92_28-52_) in the presence of increasing concentrations of 2,2,2-trifluoroethanol (TFE). **(D)** AlphaFold2 multimer model of Pho92 (cyan) and CNOT9 (limegreen), Pho92 W5 (magenta), Pho92 28-52 (magenta). See figure S2C and S2D for further details on predicted interaction surfaces and residues. (**E**) Biolayer interferometry (BLI) analysis of the Pho92 peptides (Pho92_1-25_ and Pho92_28-52_) and human Caf40/CNOT9. BLI plot shows the association and dissociation kinetics of recombinant CNOT9 to immobilised Pho92 peptides. Biotinylated peptides were loaded onto streptavidin (SA) biosensors. A representative experiment of n=3 is shown. (**F**) Similar analysis to (E), except that peptide Pho92_1-25_ with tryptophans (W5 and W18) region and the same peptide region with alanine substitutions of the tryptophan residues (Pho92 _1-25, W5A_, Pho92_1-25, W18A_, Pho92 _1-25_, _W5A/W18A_) were used. (**G**) Co-immunoprecipitation of Caf40 and Pho92. We used V5-tagged Caf40 and *CUP1* promoter-inducible HA-Pho92. For the analysis, we used HA-Pho92 in the absence and/or presence of Caf40-V5 (FW10829 and FW11775). We precipitated with Caf40-V5 from extracts treated or not treated with RNAse A. Input and eluate were assessed by immunoblotting with V5 and HA antibodies. (**H**) Similar analysis as in G, except that *pho92-W5A* and *pho92-W18A* single and double mutants were used (FW11938, FW11945, FW11971). (**I**) Similar analysis as in G, except that *pho92-*Δ*28-52* and *pho92-*Δ1-60 and *pho92-*Δ1-70 deletions in the IDR of Pho92 were tested for interaction with Caf40 (FW12128, FW12068, FW12288).

To test whether the region Pho92 (28-52) adopts α-helical structure, we used circular dichroism (CD) spectroscopy to analyze the structure of a peptide spanning residues 28 to 52 (Figure 2C)^24^. Far-UV CD spectra of the peptide in aqueous buffer are typical of a random coil conformation, suggesting that, in isolation, the peptide is largely unstructured. We then titrated 2,2,2-trifluoroethanol (TFE), a solvent that stabilises peptide secondary structure, to investigate the propensity of the peptide to adopt helical conformation. Spectra recorded with increasing TFE concentrations show a random-coil to helical transition, with maximum effect reached at 30% TFE (Figure 2C). This confirms that the peptide has a high propensity to form α-helical structure and it is thus likely to be helical when full-length Pho92 is in complex with cellular partners.

To determine whether Caf40 and Pho92 interact directly, we performed biolayer interferometry (BLI) using Pho92 peptides and purified recombinant human Caf40 (CNOT9), as yeast Caf40 could not be produced recombinantly. Given the high sequence and structural conservation between yeast Caf40 and human CNOT9, including the predicted Pho92-binding interfaces, we leveraged the human ortholog for these biophysical assays (Figures 2D, S2C and S2D). Recombinant human CNOT9 was expressed and purified from *E. coli* (Figure S2E). We first tested the binding of the Pho92-IDR tryptophan-containing and α−helical regions (Pho92_1-25_ and Pho92_28-52_, respectively) to CNOT9. We observed direct interactions with K_d_ values of 15 ± 2 μM for Pho92_1-25_ and 155 ± 21 μM for the Pho92_28-52_ peptide (Figure 2E, Table S1).

Next, we tested whether the Pho92_1-25_ peptide required tryptophan residues for its interaction with Caf40/CNOT9. We generated tryptophan-to-alanine substitutions (W5A, W18A, and W5A/W18A) (Figure 2F). Both the Pho92_1-25, W5A_ and Pho92_1-25, W18A_ single substitutions displayed a strong reduction in affinity, with K_d_ values of 400 ± 50 μM and 1220 ± 160 μM, respectively, while the double mutant (Pho92_1-25, W5A/W18A_) peptide displayed even weaker binding (K_d_ > 3000 μM; Figure 2F, Table S1). Together, these data demonstrate that tryptophan residues within the N-terminal IDR (Pho92_1-25_) mediate a relatively high-affinity interaction with Caf40/CNOT9, whereas the α-helical region (Pho92_28-52_) exhibits a lower affinity.

### In cells, Pho92 interaction with Caf40 requires the α−helical region in the IDR

To test the interaction between Caf40 and Pho92 in cells, we performed co-immunoprecipitation. We generated a strain with Caf40 tagged with V5 and Pho92 tagged with HA expressed from an inducible promoter. We found that Pho92 co-immunoprecipitated with Caf40-V5 (Figure 2G). The Pho92–Caf40 interaction was resistant to RNase treatment, indicating an RNA-independent interaction. Next, we mutated the two tryptophans (*pho92-W5A* and *pho92-W18A*). The interaction between Pho92 and Caf40 was retained in the single and double mutants (*pho92-W5A* and *pho92-W18A*) (Figure 2H). Thus, the tryptophan residues do not contribute to the interaction with Caf40 in cells. Second, we deleted regions in the N-terminal (*pho92-*Δ1-70 and *pho92-*Δ1-60) and specifically deleted the α−helical region (residues 28 to 52, defined as *pho92-*Δα) in the presence or absence of W5A/W18A substitutions (Figure 2I and 2J). All three N-terminal deletion mutants (*pho92-*Δ1-70, *pho92-*Δ1-60, and *pho92-*Δα) showed strongly reduced interaction with Caf40. While both the α−helical region and tryptophan residues can mediate Caf40 binding *in vitro*, the interaction in cells is driven primarily by the Pho92 α−helical region embedded in the IDR, suggesting that the tryptophan contacts observed *in vitro* may not be the dominant binding site in cells.

### The Pho92 IDR and predicted **α−**helical region are required for the decay function in a tethered reporter context

Having established that the Pho92 IDR engages Caf40 through its α-helical region in cells, we next asked which regions within the IDR are required for decay activity. A previous study demonstrated that the Pho92 IDR promotes decay in a heterologous tethered reporter and identified the N-terminal region as the primary functional domain^25^. Here, we extended this analysis to define the specific structural determinants required for decay activity, and critically, validate these requirements at endogenous m6A-modified transcripts during meiosis.

We first repeated the IDR analysis in a heterologous reporter. In short, we fused Pho92 fragments to the λN protein sequence, tethered them to boxB sites in a YFP reporter, and used RFP as a normaliser (Figure 3A, left panel; Figure S3A)^25^. We monitored YFP, RFP, and iRFP (Pho92 fragments) signals by flow cytometry. Consistent with previous findings, we observed an 80% reduction in reporter activity for the full-length and the N-terminal IDR fragment (residues 1 to 155) (Figure 3B, middle panel). We observed a similar decrease in YFP RNA levels for both the full-length and N-terminal IDR fragment (Figure 3B, right panel). We also examined how deletion of the α−helical region affects reporter activity. We generated a full (residues 28-52, Δα) and a smaller deletion of the predicted α-helical region (residues 35 to 42, Δαs). Consistent with interaction analysis, both the deletions (Δα and Δαs) in N-terminal IDR fragments did not repress reporter activity, suggesting that this region is required for the decay of m6A-modified transcripts (Figure 3B, middle and right panels).

**Figure 3.**
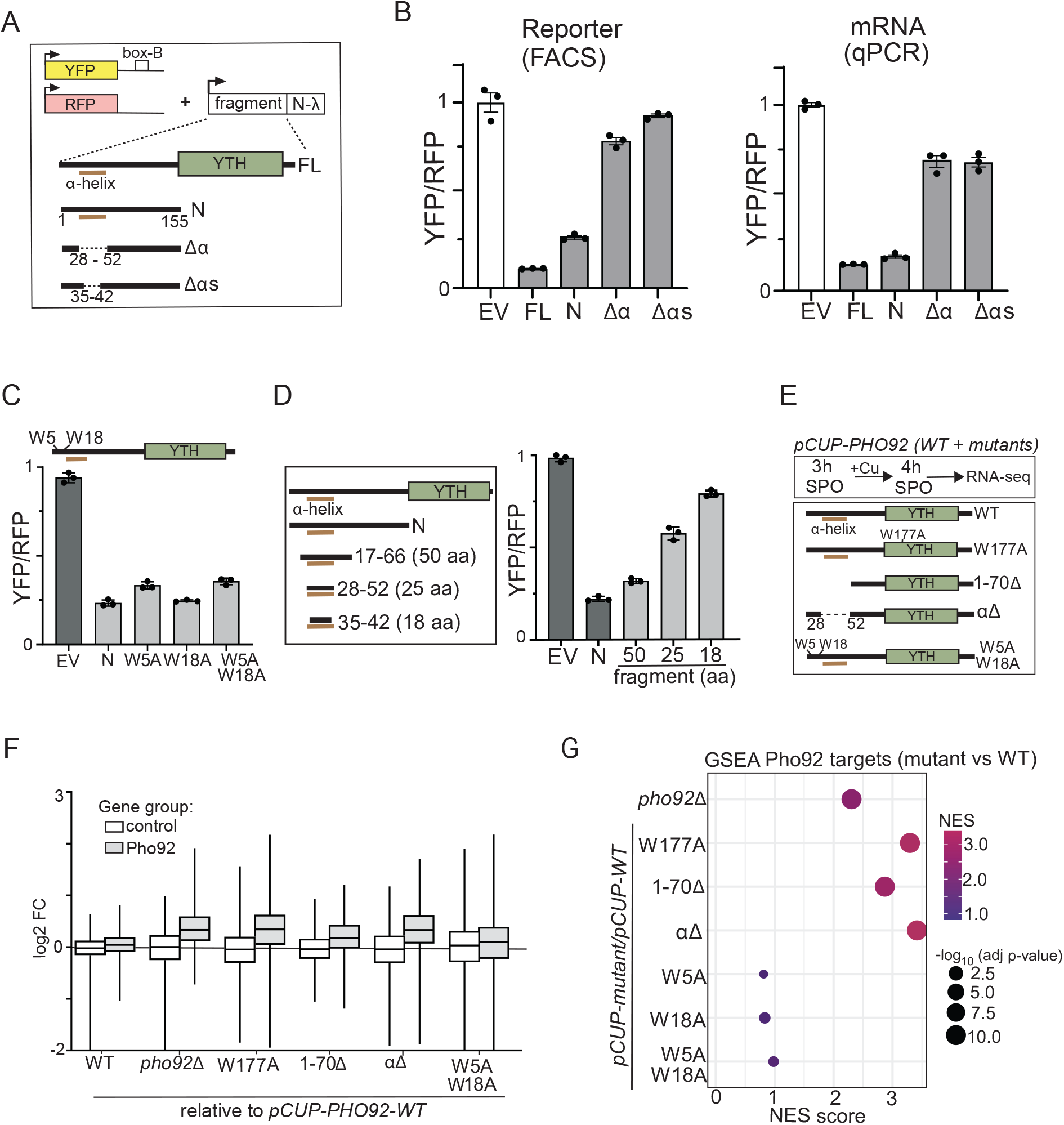
The IDR and α-helical region are essential for Pho92 effector function. (**A**) Scheme of dual fluorescence reporter described previously (left). Pho92 (and mutants) were fused to the λν protein and tethered to YFP mRNA harbouring box B sequences. RFP was used to normalise the signals. **(B)** We determined fluorescence signal by flow cytometry (FACS), and RNA levels by qPCR. Shown are empty vector control (EV), full-length Pho92 (FL), the N-terminal IDR (N, residues 1 to 155), and N-terminal IDR with the α-helical region deleted (28-52) or partly deleted (35-42) (Δα and Δαs) (FW855, FW859, FW937). The mean signal (and SD) relative to the EV control is shown across 3 biological replicates. **(C)** Same assay as in A, except assessing the function of W5A, W18A and W5A/W18A residues in the N-terminus of the Pho92 IDR (FW855, FW856, FW857 and FW858). (**D**) Same assay as in A, except assessing different-sized N-terminal fragments of the Pho92 IDR (FW847, FW857, FW848, FW967, and FW969). The mean signal (and SD) relative to the empty vector (EV) control is shown across 3 biological replicates. (**E**) Scheme of RNA-seq setup for testing IDR deletions and mutations (WT, W177A, Δ1-70, Δα and W5A/W18A) (FW10892, FW12069, FW12439 and FW11878). Pho92 was expressed from the *CUP1*-inducible promoter. We induced cells to enter meiosis (3 hours in SPO), and subsequently induced Pho92 expression for 1 hour with copper sulphate. (**F**) Box plots (with median and inter-quartile range) comparing the control set of transcripts to the Pho92 targets for the different IDR mutants described in D. (**G**) GSEA plot of Pho92 targets compared across *pho92*Δ and the different IDR mutants.

Consistent with our co-immunoprecipitation data showing that tryptophan residues are dispensable for Caf40 binding in cells, the combined substitution of W5 and W18 (W5A/W18A) did not affect reporter repression, indicating that the tryptophan residues are dispensable (Figure 3C). Finally, we determined the tethered reporter activity across different N-terminal fragments of Pho92. We found that the region encompassing residues 17 to 66 was sufficient to repress the reporter to a level almost comparable to the full-length N-terminus (70% versus 80% reduction) (Figure 3D). Smaller fragments (residues 35 to 42 and residues 28 to 52), despite containing the α-helical region, were less potent (40% and 20% reduction), suggesting that additional residues outside the predicted α-helical region contribute to full activity. Together, these results suggest that the α-helical region, rather than the tryptophan residues, is the primary functional determinant within the Pho92 IDR that is required but not sufficient for decay function.

### Pho92-IDR **α**-helical region is required for RNA decay in meiotic cells

To further test these observations for endogenous targets, we examined the expression of Pho92/m6A target transcripts by RNA-seq in meiotic cells expressing IDR deletion mutants. Previously, we showed that Pho92 associates in an m6A-dependent manner with approximately 500 transcripts^9^. Consistent with a role in mRNA destabilisation, cells entering meiosis that lack Pho92 (*pho92*Δ) display a mild increase in expression levels of Pho92 target transcripts^9,10^. Moreover, the overall turnover of m6A-modified transcripts is reduced in *pho92*Δ cells entering meiosis^9^.

We expressed Pho92 IDR mutants from the *CUP1*-inducible promoter to ensure uniform expression during meiotic entry, induced expression at 3 hours in SPO, and collected RNA-seq samples 1 hour after induction (Figure S3B). All *pho92*-IDR mutants were expressed at comparable levels, as confirmed by Western blot (Figure S3B). We leveraged the expression profile of *pho92*Δ cells to assess the RNA-seq profiles for the IDR deletion mutants. In short, for the analysis, we compared the Pho92/m6A targets to a control set of transcripts (no m6A and no Pho92 binding). As expected, *pho92*Δ cells showed a marked increase in expression of Pho92 targets (mean log2 FC of 0.26), while the control set of transcripts was not affected (mean log2 FC of 0.01) (Figure 3F). Deletion of the IDR region encompassing the helical region and tryptophan residues (*pho92-*Δ*1–70*) resulted in a significant increase in Pho92 target transcript levels relative to the control, whereas the W5A/W18A mutant (*pho92-W5A/W18A)* had little effect on Pho92 target transcript levels (Figures 3F and 3G). The effect of the α-helical region deletion (*pho92-Δ*α) was comparable in magnitude to that of *pho92-W177A*, which cannot associate with m6A-modified transcripts (Figures 3F and 3G). These data establish that the α-helical region in the IDR, but not the tryptophan residues (W5 and W18), is required for the turnover of endogenous m6A-modified transcripts.

### Caf40 is specifically required for the turnover of m6A-modified transcripts in meiosis, but not in tethered reporter

Having established that the Pho92 IDR drives m6A-transcript decay through its α-helical region, we next asked which RNA regulatory factors mediate this activity in meiotic cells. Although our biochemical and proximity labelling data implicated Caf40 as a direct Pho92 interactor, it remained unclear whether Caf40 is functionally required for m6A-transcript turnover, and whether other RNA regulatory pathways contribute. To address this in an unbiased manner, we generated depletion alleles for nine RNA regulatory factors spanning multiple decay pathways, including subunits of the Ccr4-NOT complex (Caf40, Not5, and Pop2), 5′-to-3′ exonucleases (Xrn1 and Rat1), nuclear decay (Rrp6), deadenylation (Lsm1), and decapping (Pat1 and Dcp2).

Consistent with a specific role for Caf40 in the m6A pathway, we found that loss of Caf40 also affects meiotic progression. Pho92 has a critical role in promoting the onset of meiosis and the viability of meiotic progeny. Unlike most other Ccr4-NOT subunits, Caf40 is not essential for vegetative growth, and its function in meiosis had not previously been assessed. We examined the onset of meiosis of *caf40*Δ cells and found a significant delay in meiotic divisions. About 60% completed meiosis at 24 hours in the *caf40*Δ compared to 90% in the wild-type control (Figure 4A). The delay in the onset of meiosis observed in *caf40*Δ cells was slightly more severe than that of *pho92*Δ, suggesting that loss of Caf40 somewhat exceeds the meiotic defects associated with disrupted m6A-modified transcript turnover.

**Figure 4.**
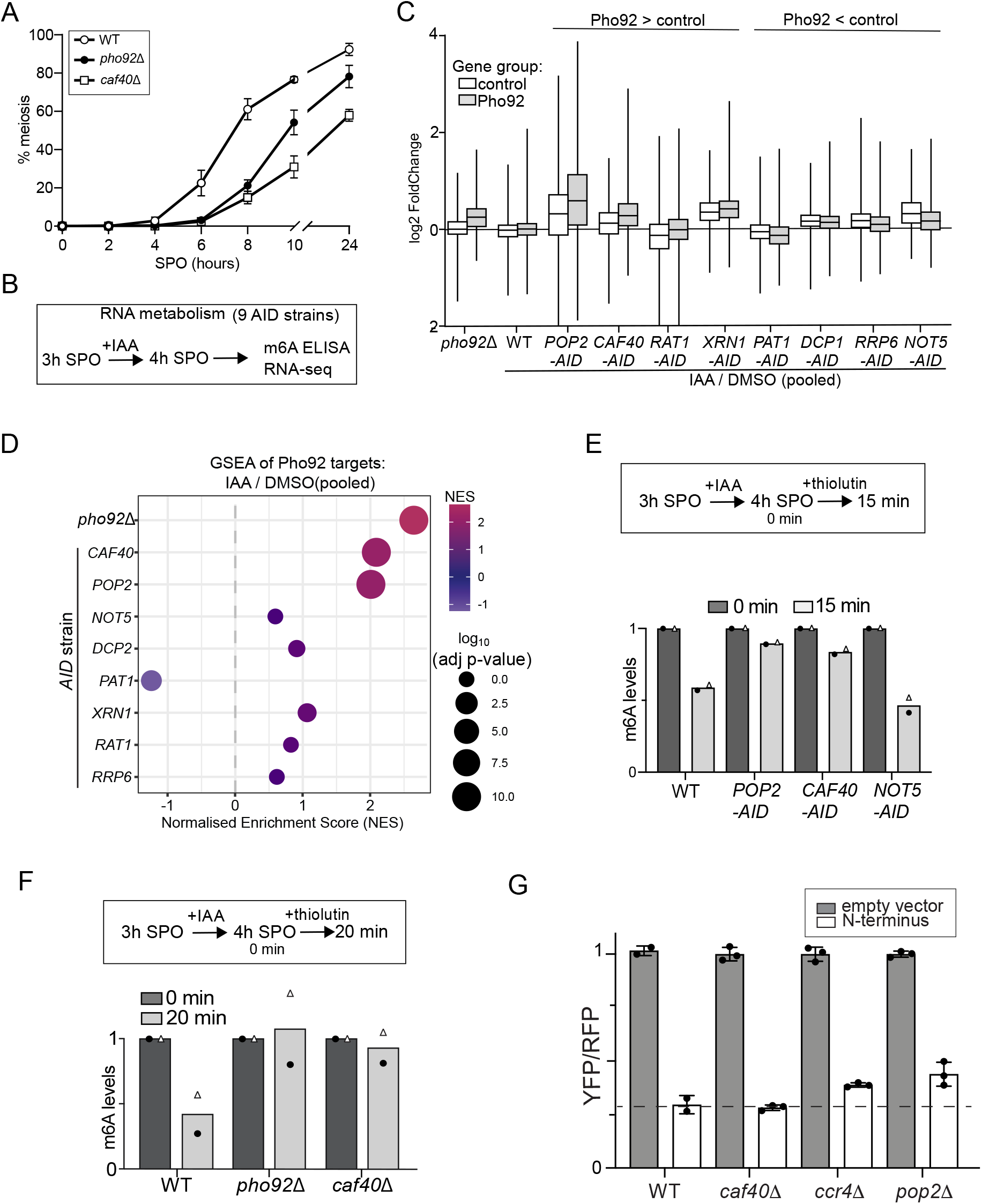
Caf40 is specifically required for the turnover of m6A-modified transcripts, but not in the tethered reporter. (**A**) Onset of meiosis as determined by DAPI counting of meiotic divisions of wild-type, *caf40*Δ, and *pho92*Δ cells. In short, cells were induced to enter meiosis, fixed at the indicated time points, and stained with DAPI. Cells with 2 or more DAPI masses were considered meiotic. At least 200 cells were counted. The mean and SEM of n=3 biological repeats are shown. **(B)** Scheme of experimental setup. We use 9 AID strains (FW5745, FW5730, FW5953, FW5958, FW6045, FW6062, FW6067, FW6072 and FW7144) harbouring the TIR1 ligase expressed from a *CUP1*-inducible promoter. Cells were induced to enter meiosis for 3 hours in sporulation medium (SPO). Subsequently, cells were treated with IAA and copper sulphate for 1 hour to deplete the RNA regulatory factors, and samples were taken for RNA-seq. As positive and negative controls, we used wild-type cells expressing the *TIR1* ligase and *pho92*Δ cells (FW5737 and FW3828). (**C**) RNA-seq analysis of RNA regulatory depletion strains. Shown are boxplots (with median and interquartile range) of Pho92 targets described in a previous study, to which Pho92 associates in an m6A-dependent manner. We also included a control set of transcripts that served as a negative control. The RNA-seq signal of the depletions (AID + IAA) was normalised to the DMSO control (pooled for all AID strains). The RNA-seq signal for *pho92*Δ was directly compared to the wild-type control. The box plots are sorted by higher or lower mean signal in Pho92 targets relative to the control. (**D**) Similar analysis described in C except that data were analysed using Gene Set Enrichment Analysis (GSEA) for the Pho92 target set of transcripts. (**E**) Relative m6A transcript stability determined by transcriptional block (+thiolutin) and chase and m6A-ELISA. In short, cells (*CAF40-AID*: FW5958, *POP2-AID*: FW5953, and *NOT5-AID*: FW6067) were induced to enter meiosis (3 hours in SPO), AID strains were treated with IAA for 1 hour and subsequently treated with thiolutin to block transcription for 15 minutes. Samples were taken at the indicated time points. The m6A signals are represented relative to the 0-minute time point. The mean of n=2 biological repeats are shown. (**F**) Similar analysis as in E, except that *caf40*Δ and *pho92*Δ cells (FW1511, FW11725 and FW3525) were used for the analysis. Cells were induced to enter meiosis at 4 hours in SPO and treated with thiolutin. The mean of n=2 biological repeats are shown.

We used the auxin-induced degron (AID) system for rapid depletion of RNA regulatory factors^26^. We induced the AID-tagged strains to enter meiosis (sporulation medium (SPO) at 3 hours) and treated the cells with Indole-3-acetic acid (IAA) to induce rapid depletion for 1 hour (Figure S4A and S4B). We also treated cells with copper sulphate to induce the *TIR* ligase controlled by the copper-inducible promoter. To ensure that m6A deposition was not affected by the rapid depletions, we measured m6A using m6A-ELISA. The nine RNA regulatory factors were depleted (AID strain treated with IAA) efficiently (Figure 4B). Except for the *LSM1-AID* strain, m6A levels were not negatively affected after depletion of the RNA regulatory factors (Figure S4B). Since reduced m6A levels in *LSM1-AID* cells would confound the interpretation of transcript stability changes, we excluded the *LSM1-AID* strain from further analysis. Notably, *POP2-*AID cells treated with IAA displayed a marked increase in m6A levels compared to the wild type, suggesting that Pop2 depletion leads to increased stability of m6A-marked transcripts (Figure S4B).

Next, we assessed the RNA-seq profiles of AID alleles. We found that Caf40 depletion (*CAF40-AID + IAA*) led to upregulation of Pho92/m6A targets compared to the control set of transcripts (Figure 4C, 4D and S4D). The mean increase of Pho92/m6A targets in Caf40-depleted cells was comparable to that of *pho92*Δ (mean log2 FC of 0.26 vs 0.37). Additionally, Pop2 depletion (*POP2-AID*) had a significant effect on Pho92/m6A targets and showed a profile comparable to Caf40 depletion (*CAF40-AID* + IAA) and *pho92*Δ mutant (mean log2 FC: 0.71 vs 0.37 vs 0.26). Not5 depletion (*NOT5-AID + IAA*) did not show a significant increase in Pho92/m6A targets and the control set of transcripts. The depletion of other RNA regulatory factors (Xrn1, Dcp2, Rrp6, Rat1, and Pan3) had little effect on the expression of Pho92/m6A targets compared to the control (Figure 4C, 4D and S4D). Thus, our data suggest that Caf40 and Pop2 subunits of the Ccr4-NOT complex regulate the stability of Pho92/m6A targets.

Since RNA-seq does not directly measure mRNA stability, we further assessed whether Caf40 and Pop2 regulate the stability of m6A-modified transcripts. To test this, we blocked transcription using thiolutin and measured m6A levels using ELISA in poly(A)-purified mRNAs, an assay we previously described (Figure 4E)^9,27,28^. Since m6A-modified transcripts have a higher decay rate than transcripts with no m6A modification, the overall m6A signal decreased (about 50%) after blocking transcription (thiolutin treatment for 15 minutes) (Figure 4E). In *pho92*Δ cells, m6A levels did not decrease upon blocking global transcription because m6A-modified mRNAs are more stable (Figure 4E). In Caf40 and Pop2-depleted cells (*CAF40-AID* and *POP2-AID + IAA*), m6A levels remained unchanged after blocking global transcription, supporting their role in promoting the degradation of m6A-modified transcripts (Figure 4E). Consistent with RNA-seq analyses, the m6A signal decreased to 50% after blocking transcription in Not5-depleted cells (*NOT5-AID + IAA*), indicating that the Not5 module of the Ccr4-NOT complex is not involved in the decay of m6A transcripts (Figure 4E). Given that *caf40*Δ cells were viable in meiosis, we repeated the transcription block m6A-ELISA assay with *caf40*Δ cells and observed no decrease in m6A signal before or after thiolutin treatment (Figure 4F).

Having established that the N-terminal IDR is sufficient for the turnover of the heterologous reporter, we next determined which Ccr4-NOT subunits mediate the activity. We tested the N-terminal IDR fragment tethered reporter in *caf40*Δ, *ccr4*Δ, and *pop2*Δ cells (Figure 4G). Surprisingly, reporter repression was retained in *caf40*Δ cells. The *ccr4*Δ and *pop2*Δ mutants showed a small partial increase in tethered IDR reporter activity compared to the wild-type control (Figure 4G). This suggests that the IDR of Pho92 can drive decay through redundant interactions with decay factors in the tethered reporter. We conclude that the Caf40 and Pop2 subunits of the Ccr4-NOT complex are specifically required for the turnover of the endogenously m6A-modified transcript in meiosis, but not required in the tethered reporter.

### Pho92 IDR is promiscuous in a heterologous reporter context

Our findings on the Pho92 IDR present a functional paradox. While interactome and AlphaFold predictions suggest binding between Pho92 and multiple Ccr4-NOT subunits (Caf40 and Not1), our *in vivo* assays showed that disrupting Caf40 alone is sufficient to abrogate the decay of endogenously m6A-modified transcripts during meiosis. We considered that this discrepancy might reflect a cellular context that restricts the IDR to a single essential interface, and that removing this context could reveal additional, redundant modes of effector interactions. To test this, we used a previously described dual-fluorescent tethered reporter that can be assayed under standard growth conditions, independent of meiosis^25^. This allowed us to ask whether Pho92 and its effectors contain capacity for plastic interactions outside their native cellular setting.

To address this, we examined whether Pho92 retains additional interactions with Ccr4-NOT in the absence of Caf40. We determined the Pho92 interactome in *caf40*Δ cells by proximity labelling using Pho92-TurboID. As expected, Pho92-TurboID was strongly enriched compared to the control (Figures 5A and 5B). Notably, Not4 and other Ccr4-NOT subunits remained enriched in *caf40*Δ cells, indicating that Pho92 retains additional contacts with the Ccr4-NOT complex independently of Caf40 (Figures 5A and 5B). Strikingly, Not1 was strongly enriched (over 8-fold) in *caf40*Δ cells, while Not1 was not significantly enriched in *CAF40* wild-type cells (Figures 5A and 5B). The Not1 enrichment in *caf40*Δ cells was comparable to Pho92 itself, indicating that both proteins are in proximity. This strongly suggests that Pho92 can interact with Not1 or other subunits of the Ccr4-NOT complex, and that Caf40 restricts Pho92 access to Not1.

**Figure 5.**
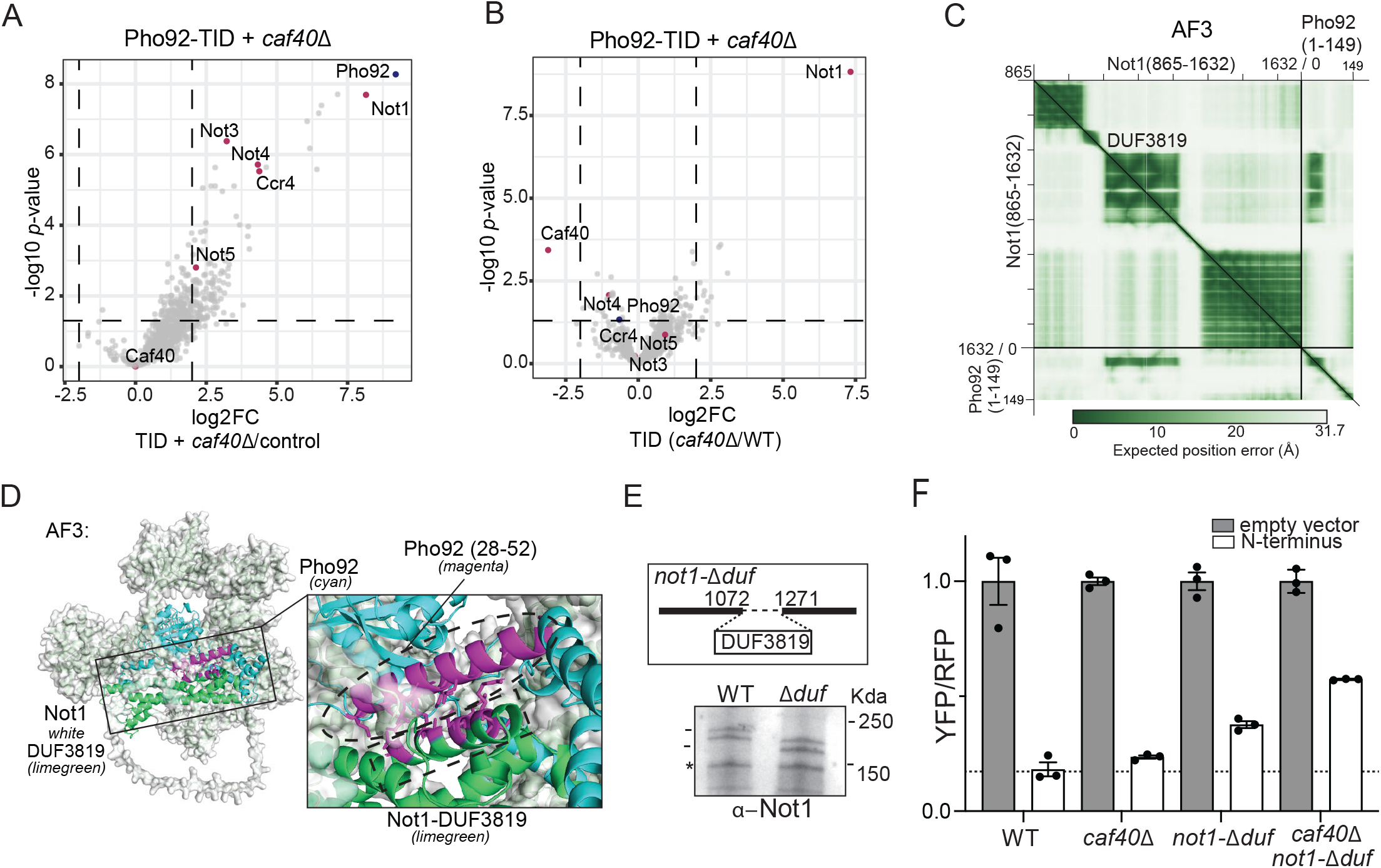
Pho92 IDR is promiscuous in a tethered reporter context. (**A**) Volcano plot showing the comparison of profiles of Pho92-TID *caf40*Δ (FW12148) to wild-type control (FW1511). (**B**) Same analysis as in A except that Pho92-TID proximity labelling profiles (wild-type *CAF40* and the *caf40*Δ (FW11105 and FW12148)) of cells were directly compared. (**C**) AlphaFold3 PAE-plot showing Not1 (residues 865-1632) highlighting the DUF3819 domain (residues 1072 to 1271) and Pho92 (residues 1-149) predicted interactions. (**D**) Representative AlphaFold3 model of Not1 (white)-Pho92 (cyan). Highlighted predicted interaction of the DUF3819 (1072-1271, limegreen) region with the α-helical region (28-52, magenta) in the Pho92 IDR. Pho92 I28, L32, L35, L38, I39, L42 residues (magenta) and Not1 R1090, V1091, Q1093, M1094, A1097, K1098, R1101, L1104, L1105 (magenta) are highlighted as consistent interaction sites across models (PAE<15, Å<5). (**F**) Reporter assay signals for the N-terminal IDR fragment in the wild-type, *caf40*Δ, *ccr4*Δ, and *pop2*Δ background cells (FW12877, FW13091, HC053, HC060). (**H**) Scheme showing CRISPR deletion of the DUF3819 domain, *not-*Δ*duf1*. Not1 expression was determined by Western blot using Not1 antibodies in wild-type and *not-*Δ*duf1* cells (FW12877 and FW13065). (**I**) Reporter assay of the N-terminal IDR fragment in wild-type (WT), *caf40*Δ, *not1-*Δ*duf* and *caf40*Δ/*not-*Δ*duf1* cells (FW12877, FW13091, FW13065, FW13091). The mean signal (and SD) relative to the empty vector (EV) control is shown across 3 biological replicates.

To explore the molecular basis for the Not1 and Pho92 interaction, we performed pairwise AlphaFold3 modelling of Pho92 and Not1 (Figures 5C, 5D and S5A-D). The AlphaFold3 analysis showed that the α-helical region in the Pho92 IDR docks into the DUF3819 domain of Not1, which is also the Caf40 binding site in Not1 (Figures 5C, 5D and S5A-D). We propose that Caf40 occludes the Pho92-Not1 interface, directing the same region in the IDR toward Caf40 as the primary effector during meiosis.

To identify redundant interfaces of the Pho92 IDR, we examined the contribution of the DUF3819 domain in Not1, the docking site for Caf40 in Not1 and the predicted binding site for the α-helical region in the Pho92 IDR (Figure 5C). We generated a deletion of the DUF3819 domain (*not1-*Δ*duf*) by CRISPR editing. The *not1-*Δ*duf* mutant did not affect cell viability, and Not1 protein levels were comparable to the wild type, allowing further analysis (Figure 5D). The *not1-*Δ*duf* significantly suppressed Pho92-IDR reporter repression (0.2 in wild-type cells and 0.4 in *not1-* Δ*duf* cells), implicating the DUF3819 domain as a redundant interface available to the tethered IDR (Figure 5E). Remarkably, when we combined *not1-*Δ*duf* + *caf40,* suppression of reporter activity was markedly increased compared to *not1-*Δ*duf* alone (0.6 in *not1-*Δ*duf* + *caf40*Δ cells versus 0.2 in wild-type cells) (Figure 5E). Taken together, these data suggest that Pho92 IDR is inherently promiscuous. When tethered to a target, it can recruit the Ccr4-NOT complex through redundant modules. Specifically, the DUF3819 domain of Not1 can independently facilitate decay via a direct interaction with the IDR. This contrasts with the regulation of endogenous m6A-modified transcripts, for which Caf40 is an essential, non-redundant requirement for decay. Thus, the Pho92 IDR selects for Caf40 specifically in the context of endogenous m6A-modified mRNAs, a selectivity that is bypassed in the heterologous reporter system.

### A single point mutation in the N-terminal IDR of Pho92 disrupts decay activity

To identify the key residues within the α-helical region of Pho92 IDR that mediate Caf40 interaction and facilitate turnover of m6A-modified transcripts (Figure S2A and S2B), we aligned the Pho92 α−helical region with other known Caf40/CNOT9-interacting proteins. CNOT4 and TTP (Human), Roquin and Bag of marbles (Bam) (*Drosophila*) have been reported or predicted to engage the concave surface of Caf40/CNOT9 via an α-helical motif (Figure 6A)^29–32^. The alignment analysis revealed enrichment of leucine residues, which possibly drive hydrophobic docking into the concave surface of Caf40/CNOT9 (Figure 6A). Indeed, leucine residues in Bam are required for the Caf40 interaction and the turnover of target transcripts.

**Figure 6.**
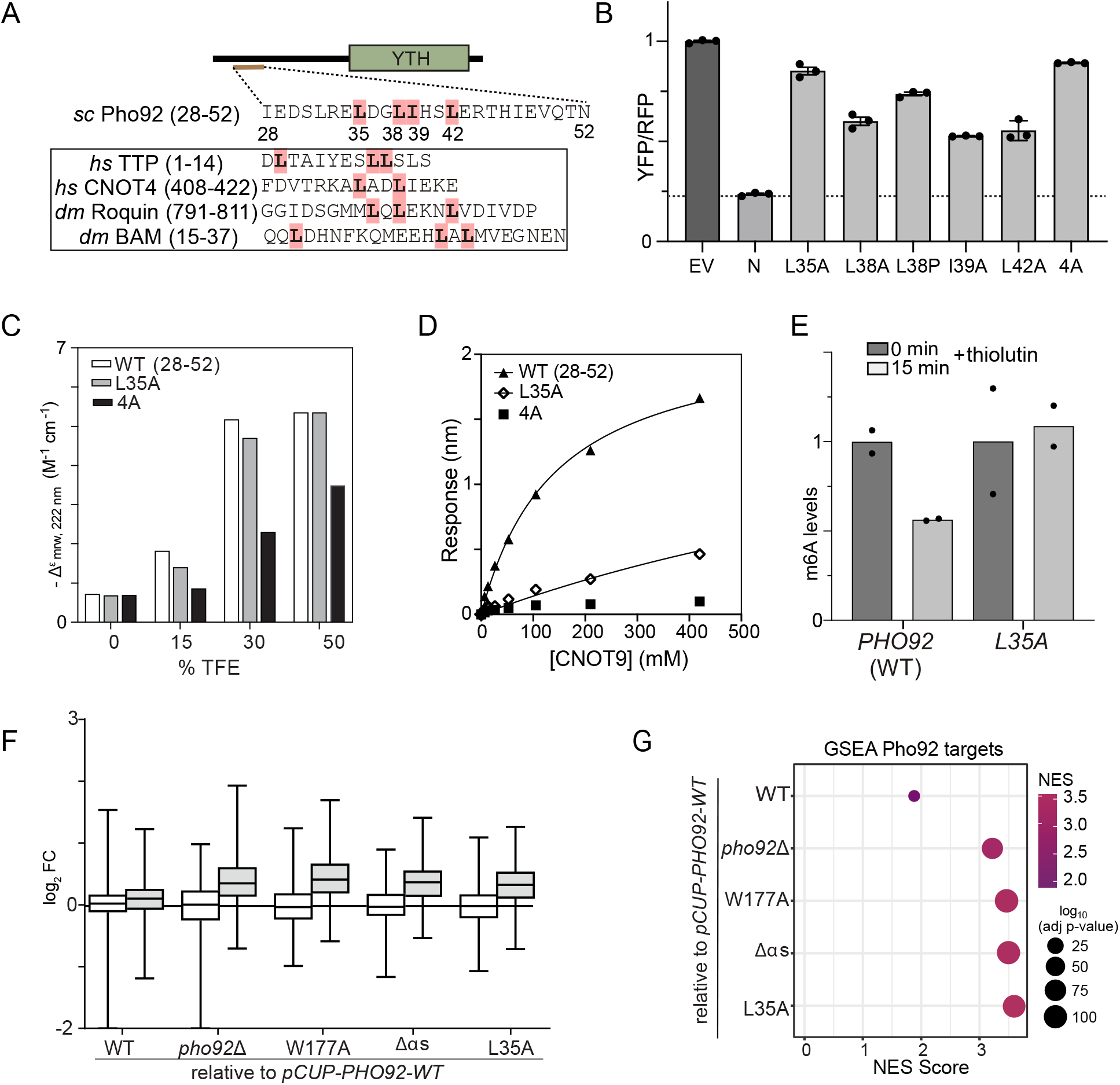
A single hydrophobic residue is essential for Pho92 effector function. (**A**) Conservation of α-helical region in Pho92 IDR comparing the sequences to *human* TTP and CNOT4, and *Drosophila* Roquin and Bag of marbles (BAM). Highlighted are the hydrophobic leucine residues. (**B**) Reporter assay of point mutations of the leucine residues in the α-helical region of the Pho92 IDR (L35A, L38A, L38P, I39A, L42A and 4A (combined alanine substitutions of L35/L38/I39/L42) (FW939, FW876, FW877, FW938, FW940, and FW941). The mean signal relative to the empty vector (EV) control is shown across 3 biological replicates. (**C**) Circular Dichroism (CD) spectroscopy of Pho92_28-52_, Pho92_28-52_, _L35A_, and Pho92_28-52_,_4A_ peptides in the presence of increasing concentrations of 2,2,2-trifluoroethanol (TFE). Shown is the negative delta signal at 222 nm. (**C**) Interaction between Pho92 IDR peptides (28 to 52 residues) Pho92_28-52_ (WT), Pho92_28-52_, _L35A_ and Pho92_28-52_,_4A_ with CNOT9 determined by BLI. (**E**) Relative m6A transcript stability determined by transcriptional block (+thiolutin) and chase and m6A-ELISA for *PHO92* and *pho92-L35A*. The mean signal of n=2 biological repeats are shown. Signals are presented relative to the 0 min time point. (FW11774, FW12912) (**F**) We assessed the *pho92-L35A* mutation. As controls, we included the WT, *pho92*Δ, *pho92-W177A,* and *pho92-* Δαs (FW10829, FW3528, FW10892, FW12912, FW12964 and). Box plots comparing the control set of transcripts to the Pho92 targets for the different IDR mutants. (**G**) GSEA plot comparing Pho92 targets of the different IDR mutants described in F.

We generated leucine-to-alanine substitutions and a leucine-to-proline substitution within the Pho92 IDR α-helical region (L35A, L38A, L38P, I39A, and L42A) and a combined mutant 4A (L35A, L38A, I39A, L42A) and tested their effect on reporter activity. All leucine substitutions derepressed reporter activity, approximating the empty vector (EV) negative control (Figure 6B). Notably, the L35A substitution produced the strongest effect. RNA levels of L35A and L38P mutants of the reporter displayed the same effect as the fluorescent reporter (Figure S6A). We conclude that single hydrophobic residues in the Pho92 IDR are key to the RNA decay activity in the heterologous reporter.

We further examined the *pho92-L35A* mutant *in vitro* and in the endogenous context during meiosis. Far-UV CD spectra measured for Pho92_28-52_ peptides (Pho92_28-52_, Pho92_28-52, L35A_ and Pho92_28-52, 4A_) showed that the effect of TFE in inducing α-helical structure formation was marginally reduced for Pho92_28-52, L35A_, compared to Pho92_28-52_, and substantially reduced for Pho92_28-52, 4A_ (Figure 6C and S6B). This indicates that the L35A mutation causes a relatively small change in the peptide’s helical propensity. In contrast, BLI measurements showed that the Pho92_28-52, L35A_ peptide displays strongly reduced binding to CNOT9 (K_d_ = 1410 ± 140 μM) compared to the wild-type Pho92_28-52_ (K_d_ = 155 ± 21 μM), while for Pho92_28-52, 4A_, binding was completely abolished (Figure 6D, Table S1). This suggests that the L35 side chain, rather than the formation of the α-helical region alone, is a key determinant of the Caf40/CNOT9 interaction *in vitro*.

We generated the *pho92-L35A* mutant induced from the *CUP1* inducible promoter in a similar set-up as described for Figure 5C. The *pho92-L35A* mutant was efficiently induced (Figure S6C). Using a transcription block assay combined with m6A-ELISA (as described in Figure 4E), *pho92-L35A* cells showed comparable m6A signals (0 and 15 minutes), indicating increased stability of m6A-modified transcripts (Figure 6E). Finally, we examined the expression of Pho92 targets by RNA-seq (as described in Figure 3F). The *pho92-L35A* mutant showed a comparable increase in Pho92 target transcripts to a degree comparable to *pho92-W177A* (Figure 6F and 6G). We conclude that a single hydrophobic residue substitution within the Pho92 IDR region is sufficient for disrupting Caf40 engagement and the turnover of m6A-modified transcripts in cells entering meiosis.

### The YTHDFs IDR are functionally active in yeast

In humans, three cytoplasmic m6A readers, (YTHDF1, YTHDF2, and YTHDF3) are orthologous to Pho92. Like Pho92, the YTHDF proteins harbour a C-terminal YTH domain and a large N-terminus of approximately 400 residues (Figure 7A). We characterized the structural features of YTHDF1-3 N-terminal regions using AlphaFold2-multimer, AlphaFold3, and disorder prediction algorithms. AlphaFold2 pLDDT scores were consistently below 50 across the N-terminal region of all three YTHDF proteins, indicating a lack of stable tertiary structure (Figure 7B, S7A and S7B). IUPred2A scores were more variable, with several stretches approaching or falling below the canonical 0.5 disorder threshold. However, unlike Pho92, the YTHDF N-termini lack a predicted α-helical region, suggesting that the mechanism of effector recruitment may have diverged despite conservation of disorder. We therefore investigated whether N-terminal regions of YTHDF1-3 retain a functionally conserved decay mechanism.

**Figure 7.**
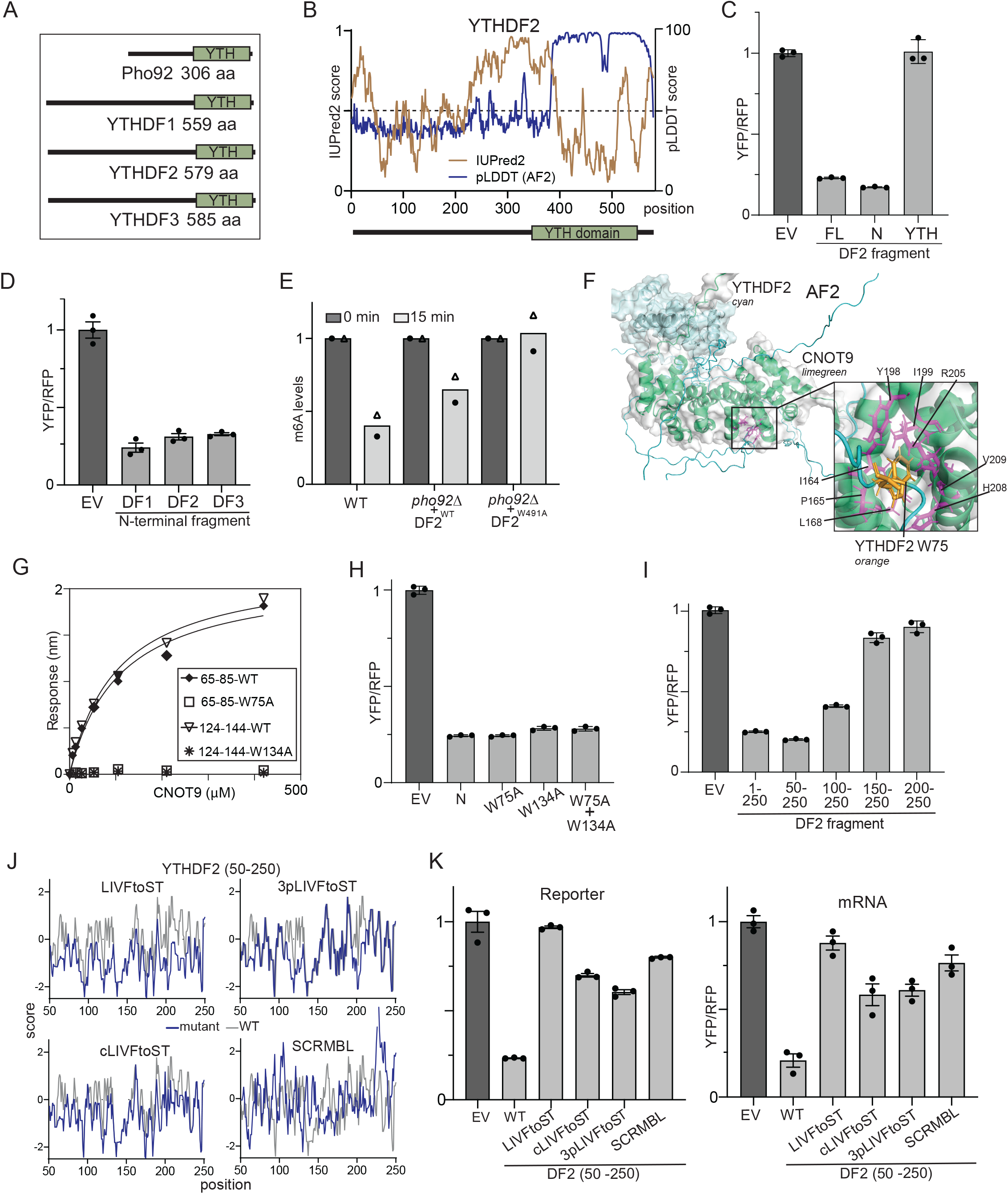
YTHDFs’ IDR hydrophobic patches are required for effector activity. **(A)** Scheme of YTHDF1, 2, and 3 compared to Pho92. (**B**) Structural disorder predictions using IUPrep2 (beige) and pLDDT (AlphaFold2) (blue) scores for YTHDF2. The x-axis indicates the residue position, and the y-axis the score. (**C**) Tethered reporter assay of full-length, N-terminal IDR of YTHDF2 and YTH domain (FW847, FW897, FW919). The mean signal (and SD) relative to the empty vector (EV) control is shown across 3 biological replicates. (**D**) The same assay as in C, except that YTHDF1-3 N-terminal fragments were compared. (**E**) Relative m6A transcript stability determined by transcriptional block (+thiolutin) and chase, and m6A-ELISA for cells expressing YTHDF2. YTHDF2 was expressed from a *CUP1*-inducible promoter in *pho92*Δ (FW10829, FW11473 and FW11991). The m6A signals are represented relative to the 0-minute time point. The mean of n=2 biological repeats are shown. (**F**) AlphaFold2-multimer interaction predictions between YTDHF2 (cyan) and CNOT9 (limegreen). Highlighted is the predicted interaction residue of YTHDF2 W75 (orange), with consistent CNOT9 predicted interacting residues I164, P165, L168, Y198, I199, R205, V209, H208 (PAE<15, Å<5) (**G**) BLI analysis of the YTHDF2 peptides (YTHDF2_65-85,_ YTHDF2_65-85, W75A,_ YTHDF2_124-144,_ YTHDF2_124-144, W134A_) and human CNOT9. BLI plot showing the association and dissociation kinetics of recombinant CNOT9 to immobilised YTHDF2 peptides. Biotinylated peptides were loaded onto streptavidin (SA) biosensors. A representative experiment of n=3 is shown. (**H**) Tethered reporter assay of YTHDF2 N-terminal fragment with tryptophan to alanine substitution (W75A, W134A and W75A/W134A). The mean signal (and SD) relative to the empty vector (EV) control is shown across 3 biological replicates. (**I**) Same assay as in H, except that different YTHDF2 N-terminal fragments were tested. (**J**) Plots of YTHDF2 N-terminal IDR hydrophobicity in wild-type and mutants according to Kyte-Doolittle hydrophobicity score^60^. We replaced hydrophobic residues (L, I, V, and F) with neutral residues (S or T) (LIVFtoST), conserved hydrophobic residues (cLIVFtoST), and 4 hydrophobic patches (4pLIVFtoST). As an additional control, we included a scrambled sequence that did not change the hydrophobicity and size (SCRMBL) (FW902, FW1085, FW1086, FW1088 and FW1090). **(K)** Tethered reporter assay of YTHDF2 N-terminal fragments described in J. The mean signal (and SD) relative to the empty vector (EV) control is shown across 3 biological replicates. (**I**) Similar analysis to H, except that RNA expression was determined by qPCR.

First, we used the tethered reporter assay to assess whether YTHDF proteins function in yeast. We fused full-length YTHDF2, the N-terminus (360 residues) and the YTH domain to the λN protein and tethered them to boxB sites in a YFP reporter. Remarkably, both the full-length and the N-terminus, but not the YTH domain alone, repressed reporter activity by more than 80%, indicating that the decay-promoting activity of YTHDF2 is mediated by the N-terminus and functionally conserved in yeast (Figure 7C). Consistent with the redundant overlapping function of YTHDFs, we found that the N-terminal IDRs of YTHDF1 and YTHDF3 also repress reporter activity to the same degree as YTHDF2 (Figure 7D).

We next asked whether YTHDF2 could functionally substitute for Pho92 in promoting m6A-mRNA decay during yeast meiosis. We expressed YTHDF2 from the *CUP1* promoter during early meiosis in a *pho92*Δ strain and monitored m6A-transcript turnover using a transcription block assay (Figure 7E). YTHDF2 expression significantly accelerated m6A-modified transcript decay, reducing m6A levels within 15 minutes of thiolutin treatment, demonstrating that YTHDF2 can functionally substitute for Pho92. The rescue depended on m6A binding, as a YTH domain mutation (W491A) abolished the effect (Figure 7E).

### The YTHDF N-terminus hydrophobicity is required for effector function

Having established functional conservation, we examined whether YTHDF2 contains similar effector interfaces in its N-terminal IDR. Inspection of the YTHDF2 IDR revealed a tryptophan motif (W75, WST motif) similar to Pho92 (which also has a WST motif) and a second tryptophan (W134). AlphaFold2 and AlphaFold3 models predicted interactions between the tryptophan-containing regions of YTHDF2 (W75 and W134) and the convex surface of CNOT9 and Caf40 (Figures 7F and S7C–I), prompting us to test these interactions biochemically. We found that two YTHDF2 peptides (YTHDF2_65-85_, YTHDF2_124-144_) spanning the tryptophan-containing regions (W75 and W134) can both directly bind CNOT9, with affinities of 104 ± 10 μM and 107 ± 12 μM, respectively (Figure 7G). The interaction was completely disrupted by alanine substitutions (YTHDF2_65-85, W75A_ and YTHDF2_124-144, W134A_). These data demonstrate that tryptophan residues in the YTHDF2 IDR can directly contact CNOT9 in vitro, though as with Pho92, these interactions are dispensable for decay activity in the tethered reporter assay (Figure 7H).

We further mapped the repressive activity within the YTHDF2 N-terminal IDR using the tethered reporter assay. We found that the N-terminal region spanning residues 1 to 250 repressed reporter activity by 5-fold, comparable to full-length YTHDF2 (Figure 7I and 7A). An IDR fragment 50 to 250 was still sufficient for reporter repression comparable to the 250-residue fragment (Figure 7I). The shorter fragments (residues 100 to 250 and 150 to 250) showed reduced repression of the reporter (Figure 7I). To determine whether repressive activity within the 50 to 150 region depends on a single defined sequence, we introduced systematic five-amino-acid linker substitutions across this region (Figure S8A). None of the individual substitutions derepressed reporter activity, indicating that no single sequence element is essential and that repressive function is distributed across multiple regions of the YTHDF2 IDR.

Our findings on Pho92 suggest that hydrophobic residues are key for interaction with Caf40 and for the repressive effector function. The N-terminal region (residues 50– 250 of YTHDF2) contains 34% hydrophobic residues (I, V, L, F, M, A, W, and C) and has a mean Kyte-Doolittle hydrophobicity of −0.2, indicating an overall hydrophilic character (Figure 7J). To test the function of hydrophobic residues in YTHDF2, we generated a panel of N-terminal mutants (Figure 7J). We generated a mutant with all the leucine, isoleucine, valine, and phenylalanine residues changed to non-hydrophobic residues (serine or threonine) (LIVF to S or T, LIVFtoST) (Figure 7J and Table S2). Moreover, we generated a construct with only the conserved residues (YTHDF1-3) mutated (cLIVFtoST), and a construct with 3 hydrophobic patches mutated (3pLIVFtoST) (Table S2 and Figure 7J). Finally, we generated a scrambled sequence (SCRMBL) with equivalent hydrophobicity to the wild-type N-terminal IDR. Notably, the mutants (except for the SCRMBL) displayed increased propensity for IDR sequence stretches compared to WT (Figure S8B).

We found that LIVFtoST completely derepressed the reporter activity, and that cLIVFtoST and 3pLIVFtoST partially derepressed reporter activity (to ∼0.6 relative fluorescence, compared to ∼0.2 for wild-type) (Figure 7K, left panel). Notably, the SCRMBL failed to repress reporter activity. To verify that reporter repression was mediated at the RNA level, we quantified RNA abundance for the same set of mutants and found that it closely tracked reporter activity (Figure 7K, right panel). This suggests that, like Pho92, hydrophobic residues in the YTHDF2 N-terminal IDR are required for the effector function. However, multiple hydrophobic patches contribute, not a single site, as we found for Pho92. The YTHDF2 N-terminal IDR strictly depends on a subset of hydrophobic residues, distributed across patches, for effector function, suggesting that the hydrophobic content within a disordered region is functionally decisive. We propose that multiple weak hydrophobic contacts can substitute for the single interface used by Pho92, while achieving a comparable functional outcome in the reporter assay.

Together, our results demonstrate that the YTHDF2 N-terminus harbours a functionally conserved effector domain within its IDR, sufficient to promote decay when tethered to RNA. As with Pho92, this domain engages the decay machinery through hydrophobic residues, indicating that IDR-mediated effector engagement is a regulatory feature conserved across more than one billion years of eukaryotic evolution.

## Discussion

How m6A reader proteins selectively recruit effector complexes to drive mRNA decay is an open question in RNA biology. Here, we identified a direct and surprisingly constrained mechanism by which the YTH domain m6A reader Pho92 promotes the turnover of m6A-modified transcripts during yeast meiosis. A short region within the Pho92 IDR engages the Caf40 subunit of the Ccr4–NOT deadenylase complex, and a single hydrophobic residue substitution is sufficient to abolish Caf40 binding and consequently m6A-transcript decay during meiosis. These conclusions have broader implications for how IDR-containing RNA-binding proteins are functionally interpreted in the cell.

A prevailing model is that intrinsically disordered regions of RNA-binding proteins drive mRNA decay through multivalent, low-affinity interactions^33–35^. Indeed, multiple biochemical studies have shown how IDRs of RNA-binding proteins can engage with multiple interaction surfaces of the Ccr4-NOT complex^32,36–38^. Our analyses of the Pho92-IDR suggest that biochemical interactions or the tethered reporter do not fully recapitulate functional output. While the Pho92 IDR is capable of engaging with multiple sites within the Ccr4–NOT subunits *in vitro* and in heterologous reporter contexts, only the Caf40 interface is functionally essential for the decay of endogenous m6A-modified transcripts during meiosis. The interaction is mediated by a short α-helical region within the Pho92 IDR that engages Caf40 directly. However, outside its native cellular context, this interface can engage additional Ccr4–NOT surfaces promiscuously.

A key insight from this work is that biochemical binding capacity, even at physiologically plausible affinities, does not predict functional contribution *in vivo*. The W5 and W18 residues of Pho92 bind Caf40 *in vitro* with affinities comparable to functionally validated interactions for other RNA-binding proteins, yet neither contributes to endogenous transcript decay^39^. This dissociation between binding and function likely reflects competition for overlapping binding surfaces *in vivo*, compounded by the requirement for active translation and cells staged in meiosis, factors that together enforce Caf40 as the dominant effector interface^9^. How active translation couples to the Pho92-Caf40-dependent decay remains an important open question. Similarly, the tethered reporter assay can give outcomes different to the physiological context. Unlike endogenous Pho92, the tethered reporter bypasses the translational requirement for decay and operates under vegetative growth conditions. The crowded environment of translating ribosomes and the relative stability of the boxB–λN tether compared to YTH domain-mediated m6A mRNA association may therefore substantially alter which contacts are accessible and productive^40^. These observations indicate that the functional output of IDR-mediated interactions is determined not by intrinsic affinity alone, but by the ensemble of competing interactions and contextual constraints present in the cell.

The mechanism described for Pho92 here has striking parallels with the *Drosophila* differentiation factor Bam. Both function as post-transcriptional regulators of gametogenesis^41,42^. Bam is an RNA-binding protein that acts in a germline differentiation context, promoting cystoblast differentiation in the Drosophila ovary and driving spermogenesis^41,42^. Like Pho92, Bam engages Caf40 via a short N-terminal α-helical region that is functionally essential for the male germline ^29,43^. The Caf40 anchor is also used in a broader context; Roquin and CNOT4 interact with CNOT9/Caf40 through hydrophobic motifs docking onto the same convex surface^30,31^. The recurrence of this docking mode across phylogenetically distant organisms, and its specific deployment in germline and gametogenic contexts in both yeast and flies, suggests that Caf40 serves as a conserved effector anchor for post-transcriptional programmes governing cell differentiation. Our identification of leucine residues in the Pho92 α-helical region as essential for Caf40 binding extends this mechanism to m6A-dependent mRNA decay in yeast meiosis.

Our findings that human YTHDF2 can functionally substitute for Pho92 and that YTHDF IDRs require hydrophobic residues for decay activity suggest that this mechanism is at least partially conserved. However, the YTHDF IDRs appear to stimulate RNA decay through distributed hydrophobic patches rather than a single dominant motif. This more distributed architecture may reflect the absence of a single dominant cellular context in human cells equivalent to yeast meiosis, resulting in a more redundant effector interface. However, it is notable that YTHDF2 decay function in human cells is also linked to active translation, suggesting that translational context may impose analogous constraints on IDR-effector selectivity in mammals^11,12^. Multiple effector interactions have been reported for YTHDF proteins *in vitro*, including YTHDF2–CNOT1 and a YTHDF3 motif essential for Pan2/Pan3-mediated deadenylation, consistent with a more promiscuous interaction landscape in human cells^4,44,45^. Whether translational coupling imposes analogous restrictions on YTHDF IDR promiscuity in mammals will be an important question for future work.

The YTHDF proteins are dysregulated in multiple cancers, and YTHDF2 in particular has emerged as a promising therapeutic target^46–49^. Our finding that hydrophobic residues distributed across the YTHDF2 IDR are required for decay activity suggests that disrupting these contacts, for example through small molecules targeting the IDR-effector interface, could represent a tractable therapeutic strategy.

More broadly, our results caution against inferring functional specificity from binding capacity alone. IDRs are increasingly recognised as hubs of protein–protein interaction, but the biological relevance of any given contact depends on whether it is permitted or enforced by the cellular environment. Defining the rules that govern this selectivity will be essential for understanding how post-transcriptional regulatory networks are organised and how their dysfunction contributes to disease.

## Methods and materials

### Yeast and plasmid construction

All strains were derived from the *Saccharomyces cerevisiae* SK1 strain background for all the meiosis experiments and the BY4741 strain background for the dual tethered reporter assay. The TurboID plasmid and pFA6a-3V5-IAA7-KanMx6 plasmid containing the 3xV5 and AID were used for C-terminal tagging^14^. We used CRISPR editing to make a gene *duf*Δ in *NOT1*^50^. In short, guides targeting the DUF3819 region in *NOT1* were introduced by Golden Gate cloning into a sgRNA vector. Donor DNA, containing homologous regions to the flanking regions of DUF3819 was ordered (Integrated DNA Technologies). The sgRNA vector was linearised with EcoRV. The Cas9 vector containing *URA3* was linearised with BsmBI. All three components were co-transformed. The genotypes of strains used are described in Table S3.

We used a *CUP1* promoter fusion *PHO92* single-copy integration plasmid described previously for generating and testing *pho92* mutations^9^. The dual (YFP/RFP) reporter system of BoxB and λN plasmids was described previously^25^. *PHO92* and *YTHDF* fragments were cloned by Gibson cloning. Site-directed mutagenesis was used to make point mutations. Synthetic fragments (IDTA) were cloned. A list of plasmids is described in Table S4.

### Growth conditions

Cells were arrested in early meiosis as described previously^51^. Briefly, yeast cells were inoculated in yeast extract peptone dextrose (YPD) media [1% weight/volume (w/v) yeast extract, 2% w/v bacto peptone, 2% w/v glucose, uracil (24 mg/l) and adenine (12 mg/l)] for approximately 24 hours, before being diluted to OD600 0.4 in buffered, yeast extract, tryptone, acetate (BYTA, (1.0% w/v yeast extract, 2.0% w/v peptone, 1% potassium acetate, 50 mM potassium phthalate monobasic) and incubated for 16-18 hours. OD600 1.8 of yeast cells were then washed and transferred to SPO (0.3% Potassium acetate, Acetic acid to pH 7.0, 0.02% D-(+)-Raffinose pentahydrate). 2 hours after transfer to SPO, CuSO4 (50 μM) and IAA (500 μM) were added to induce synchronous meiotic entry and auxin-induced protein depletion. In all steps, yeast was grown at 30°C at 300 rpm, and a 1:10 culture volume to flask total volume ratio was maintained.

### TurboID

180 OD units were washed three times with sterile water, then with 25 mL of 100% acetone, dried and resuspended in 5 mL TE buffer (100 mM Tris pH 7.5, 1 mM EDTA, 2.7 mM DTT), per 18 OD units.) 500 μl of zirconia/silicate beads (Ø 0.5 mm) were added and cells were lysed in the bead-beater. 3 x SDS sample buffer (loading buffer with Βromophenol blue removed) (187.5 mM Tris pH 6.8, 30% glycerol, 3% SDS) was added and the samples were boiled at 100°C for 5 minutes. For affinity purification, 400μl of streptavidin (M-280, Thermo Fisher Scientific or S1420S, New England BioLabs) were used per sample, incubated at 4°C for 3 hours. Beads were washed once with wash buffer (50 mM Tris HCl pH 7.5, 2% SDS), four times with RIPA 0.4% SDS buffer (50 mM Tris HCl 7.5, 150 mM NaCl, 1.5 mM MgCl_2_, 1 mM EGTA, 0.1% SDS, 1% NP-40) and five times with 20 mM ammonium bicarbonate, to ensure no detergent remained. Beads were resuspended in 50 μl of 50 mM ammonium bicarbonate.

### Proteomic

Beads were digested using a trypsin-based protocol prior to peptide identification by mass spectrometry. Samples were reduced with 5 mM DTT for 1 h at 37 °C in a thermomixer. Proteins were then alkylated with 10 mM IAA for 30 minutes in the dark at room temperature. After reduction/alkylation, Beads were trypsinised to digest biotinylated proteins (0.4 μg/μl trypsin in 50 mM NH_4_HCO_3_). Digestion was performed overnight at 37 °C in a thermomixer at 450 rpm. The reaction was stopped with 10% formic acid (FA). Peptides were recovered and the beads were discarded using a magnetic rack. Finally, a C18 clean-up was performed using EV2018 EVOTIP PURE. The desalting was done according to the manufacturer’s protocol. Briefly, the Evotips were conditioned with 0.1% FA in acetonitrile and equilibrated with 0.1% FA in water. Approximately 1 µg of each digested sample was loaded onto Evotips. Peptides were eluted from the Evotips with 50% acetonitrile into a vial and vacuum dried by SpeedVac to remove any traces of organic solvents. Finally, the dried peptides were resuspended in 0.1% FA.

The resulting peptides were analysed by nano-scale capillary LC-MS/MS using an Ultimate U3000 HPLC (ThermoScientific Dionex, San Jose, USA) to deliver a flow of approximately 300 nL/min. A C18 Acclaim PepMap100 5 µm, 75 µm x 20 mm nanoViper (ThermoScientific Dionex, San Jose, USA), was used to trap the peptides prior to separation on an EASY-Spray PepMap RSLC 2 µm, 100 Å, 75 µm x 500 mm nanoViper column (ThermoScientific Dionex, San Jose, USA). Peptides were eluted with a 90 min gradient of acetonitrile (2%v/v to 80%v/v). The analytical column outlet was directly interfaced via a nano-flow electrospray ionisation source to a hybrid quadrupole orbitrap mass spectrometer (Lumos Tribrid Orbitrap mass spectrometer, ThermoScientific, San Jose, USA). Data dependent analysis was carried out, using a resolution of 120 000 for the full MS spectrum, followed by MS/MS spectra acquisition in the linear ion trap using “TopS” mode. MS spectra were collected over a m/z range of 300-1800. MS/MS scans were collected using a threshold energy of 32% for collision-induced dissociation.

LC-MS/MS raw files were processed in MaxQuant (version 2.0.3.1). The LFQ algorithm and match between runs settings were selected. Data were searched against the reviewed UniProt *Saccharomyces cerevisiae* proteome using the Andromeda search engine embedded in MaxQuant. Trypsin was set as the digestion enzyme (cleavage at the C-terminal side of lysine and arginine amino acid residues unless proline is present on the carboxyl side of the cleavage site) and a maximum of two missed cleavages were allowed. Cysteine carbamidomethylation was set as a fixed modification, while oxidation of methionine and acetylation of protein N-termini were set as variable modifications. The “match between runs” feature was used with a matching time window of 0.7 min and an alignment time window of 20 min. Label-free quantification was performed using the MaxLFQ feature included in MaxQuant utilising default LFQ parameters. Minimum peptide length was set at 7 amino acid residues. FDR, determined by searching a reverse sequence database, of 0.01 was used at both protein and peptide level. The MaxQuant protein groups output file was imported into Perseus software (version 1.4.0.2) for further statistical analysis and data visualisation. Contaminant and reverse protein hits were removed. LFQ intensities were log_2_-transformed. Missing values (NaN) were inserted from a normal distribution with default values. For each sample, triplicates were grouped. Data was filtered for at least two out of three replicates LFQ intensity values in at least one group. Protein LFQ intensities were normalised and missing values were imputed by values simulating noise around the detection limit using the default parameters. A protein was considered significantly differentially expressed when FDR <0.05.

### RNA isolation

RNA was extracted from yeast as previously detailed^27^. 24 OD units were collected by centrifugation, and snap-frozen in liquid nitrogen. Per 10 OD units, RNA was extracted with Tris-EDTA-SDS (TES) buffer (10 mM Tris-HCl pH 7.5, 10 mM EDTA, 0.5% SDS) and Acid Phenol:Chloroform:Isoamyl alcohol (125:24:1, Ambion) at 65°C for 45 minutes. After centrifugation at 4°C, max speed, 10 minutes, the aqueous phase was transferred to cold ethanol with 0.3 M sodium acetate. Precipitation was carried out overnight at 4°C. rDNase (Macherey-Nagel) treatment was carried out for 20 minutes at 37°C, before spin column purification (Macherey-Nagel).

### M^6^A-ELISA

m6A ELISA was described previously^9,27,28^. In short, two rounds of oligo (dT)_25_ selection were performed to reduce ribosomal RNA (rRNA) contamination to <1%. From yeast total RNA, a minimum of 50 μg of DNase-treated and column-purified RNA was added to 900 μL of binding buffer (Tris-HCL 20 mM pH 7.4, Lithium Chloride 500 mM, Lithium dodecyl sulphate 0.5%, EDTA 2 mM) and applied to 50 μL of oligo(dT)_25_ beads (New England BioLabs, S1419S) pre-equilibrated in binding buffer. After 15 minutes of rotation at room temperature, samples were incubated on ice for 2 minutes, followed by two washes in wash buffer A (Tris-HCL 20 mM pH 7.4, Lithium Chloride 500 mM, Lithium dodecyl sulphate 0.2%, EDTA 2 mM), two washes in wash buffer B (Tris-HCL 20 mM pH 7.4, Lithium chloride 500 mM, EDTA 2 mM) ending in one wash with 10 mM Tris-HCL pH 7.4. The sample was heat eluted in 55 μL 10 mM Tris-HCL pH 7.4 after 5 minutes of incubation at 75-80°C, shaking in a thermoblock at 1400rpm. In the second round, heat elution was done in 25 μL of 10 mM Tris-HCL pH 7.4. The sample yield can be quantified by spectrophotometer (i.e. NanoDrop, Thermofisher Fisher Scientific); however for applications such as in m^6^A ELISA, quantification was performed with Qubit^TM^ RNA High Sensitivity kit (Thermo Fisher Scientific, Q32852) to improve accuracy.

Standards used for the m^6^A ELISA were generated using the MEGAscript® T7 Transcription Kit (Invitrogen, AM1334) according to the manufacturer’s instructions, with the control template included in the kit used to produce both unmodified and modified Adenosine. For generating m^6^A RNA standards, ATP was replaced with an equivalent concentration of *N*^6^-ATP (Jena Bioscience, NU-805-BIO). The generated standards were subsequently TURBO^TM^-DNase treated to remove the template DNA and column-purified (as previously described section **Error! Reference source not found.**)

90 μL of binding solution (Abcam, ab156917) was added to a clear DNA binding microplate (Abcam, ab210903). 50 ng of mRNA samples were added in triplicate. Each m^6^A-ELISA was validated by including a wild-type sample and an *ime4*Δ sample as positive and negative controls. Previously generated *in vitro* RNA samples were plated as standards. After mixing the mRNA samples into the binding solution, the plate was incubated at 37°C for 2 hours. Each well was washed 4 times with phosphate buffered saline (PBS) supplemented with 0.1% Tween-20. The sample wells were incubated with 100 μL of primary antibody solution (1:10 000 m^6^A primary antibody, ABClonal A19841, 0.5 μg/mL *ime4*Δ yeast total RNA in PBS-T 0.1%) for 1 hour RT. Each well was washed four times with PBS-T 0.1%. The sample wells were incubated with secondary antibody solution (1: 5000 anti-Rabbit Goat IgG (HRP), ab205718 in PBS-T 0.1%) for 30 minutes RT. Each well was washed five times with PBS-T 0.1%. 100 μL TMB ELISA Substrate (Fast Kinetic Rate) (Abcam, ab171524) was added to each sample well for up to 30 minutes to develop the plate. The reaction was stopped by adding 100 μL of stop solution (Abcam, ab171529). Absorbance was read out at 450 nm using a Tecan Spark® microplate reader.

### RNA-seq

RNA was extracted, DNase-treated and column purified as previously described^15^. For degron strains, libraries from 100 ng of RNA were generated using the NEBNext Ultra II Directional PolyA mRNA Kit (modules K0105 and K0078), following the manufacturer’s protocols. Paired-end sequencing (100 bp) was performed on an Illumina NovaSeq X platform, with a minimum depth of 20 million reads per sample.

For RNA sequencing of *pho92* mutant strains, mRNA enrichment was performed using the NEBNext Poly(A) mRNA Magnetic Isolation Module (NEB #7490), followed by library construction with the NEBNext Ultra II Directional RNA Library Prep Kit (NEB #E7760/E7765), according to the manufacturer’s protocols. Paired-end sequencing (100 bp) was performed on an Illumina NovaSeq 6000 platform, with a minimum depth of 20 million reads per sample.

### RT-qPCR

Reverse transcription was performed on 500ng of DNase-treated and column-purified RNA using the ProtoScript II First Strand cDNA Synthesis Kit (New England BioLabs, E6560S) according to the manufacturer’s instructions. In most cases random primer mix was used, in some applications oligo (dT) priming was used.

cDNA used as the qPCR template was diluted 1:5 with sterile water. Each qPCR reaction was made up to 10 μL (5 μL 2 x Power Up SYBR Green Master Mix (Thermo Fisher Scientific, A25742, c_f_ = 1x, 0.25 μL 10 μM forward and reverse primer (c_f_ = 250 nM) and 1 μL of the previously 5-fold diluted cDNA template)). Real-time quantification was performed on a QuantStudio^TM^ 5 or QuantStudio^TM^ 7. The cycling conditions were as follows: denaturation at 95°C for 30 seconds, followed by 40-cycles of denaturation at 95°C for 1 second, annealing and extension at 60°C for 30 seconds.

A standard curve – formed from 5 x 10-fold serial dilutions of a pool of total cDNA was used to obtain relative quantification of each primer pair and target gene. A linear plot was generated from threshold cycle values (Ct values) of each dilution for each gene. This served as a calibration curve from which the extrapolation of the relative expression of each gene at each time point is calculated. The relative expression of each gene was normalised to that of the housekeeping gene *ACT1.* The following primers were used for YFP (5’ – TCCATGGCCAACCTTAGTCAC-3’, 3’ – GAACATAACCTTCTGGCATGGC – 5’), RFP (5’ – ACAGACGGCCAAGCTAAAAG – 3’, 3’ – TGCCGGATGCTTAGTGAAAG – 5’) and *ACT1 (*5’ *–* GTACCACCATGTTCCCAGGTATT – 3’, 3’ – AGATGGACCACTTTCGTCGT – 5’).

### Flow cytometry and Heterologous reporter analysis

Three transformants were grown overnight in synthetic dropout medium and diluted to OD_600_=0.05 the next morning. They were grown for ∼3.5-5 hours to exponential phase (OD_600_=0.4-0.7). A minimum 2.5 OD units were collected, pelleted and washed with PBS. Cells were fixed in 4% paraformaldehyde (Thermo Scientific Chemicals, 047392.9M) in PBS at room temperature (RT) for 15-60 minutes in the dark. Cells were washed and resuspended in PBS before storage at 4°C. Cells were transferred to polystyrene tubes. Flow cytometry was performed on either a BD LSRFortessa or a CytoFLEX LX analyser, measuring fluorescence from YFP, RFP and emiRFP670. On the BD LSRFortessa, YFP and RFP fluorescence were measured following excitation by the 488-nm blue and 561-nm yellow-green lasers, respectively, while emiRFP670 fluorescence was excited by the 633-nm red laser. On the CytoFLEX LX, YFP fluorescence was collected using the blue 525-nm channel, RFP fluorescence using the yellow 610-nm channel, and emiRFP670 fluorescence using the red 660-nm channel. For each sample, a minimum of 10,000 cells were analysed after gating on forward and side scatter using both height and area parameters. Flow cytometry data were collected using BD FACSDiva or CytExpert software, depending on the analyser used.

### Flow cytometry analysis

Data were analysed with flowCore in R after applying gating schemes on forward and side scatter and selecting for single cells. This captures ∼60% of events from flow cytometry. The YFP:RFP fluorescence ratio is calculated for each cell and reported as a mean YFP:RFP ratio. This ratio is normalised to the highest YFP:RFP ratio (i.e. the empty vector control). Three biological replicates have been performed for the control and each test fragment, and a standard deviation of the normalised YFP:RFP ratios from the three biological replicates has been reported

### DAPI counting

At set time points following transfer of yeast cells to SPO, samples were collected by centrifugation and fixed in 80% ethanol. Samples were then resuspended in 4’,6-diamidino-2-phenylindole (DAPI) (1 µg/ml in phosphate-buffered saline, PBS). The number of cells (n = 200) that had undergone one or two meiotic divisions (and so contained 2-4 nuclei) was assessed microscopically.

### Recombinant protein expression and purification

Recombinant CNOT9 was expressed using *E.coli* BL21(DE3) competent cells (New England Biolabs, C2527H). A starter LB culture of 100 mL with 100 μg/mL Ampicillin was inoculated overnight. The next day, 5 mL of saturated culture was used to inoculate 1 L Zym 5052 with 100 μg/mL ampicillin in a baffled flask. Cells were grown to saturation at 37 °C for 11 hours, then changed to 20 °C for a minimum of 24 hours or up to 60 hours. Cells were harvested by centrifugation at 4,500 rpm, 4 °C for a minimum 10 minutes to remove the media.

Cell pellets from 1L culture were resuspended in 50 mL lysis buffer (20 mM Tris pH 7.9, 250 mM NaCl, 5% glycerol, TCEP 0.5mM, supplemented with cOmplete^TM^ Protease Inhibitor Cocktail (1 tablet per 50 mL buffer), Sigma Aldrich 11697498001) and treated with 2 μL Benzonase® (Sigma-Aldrich, E8263) for a minimum of 30 minutes rolling at 4°C. Cells were lysed with a French Press Cell disruptor (Continuous Flow Cell disruptor, Constant Systems CF2) at 4°C, 25-30 psi, in lysis buffer. Lysed cells were centrifuged at 30.000 rpm for 30 minutes at 4 °C. The supernatant (clarified lysate) was separated from cell debris and filtered through a 0.25 μM filter.

10xHIS-GST-CNOT9(16-284) was separated from the whole lysate using Glutathione Sepharose^TM^ 4B (Cytiva, 17075605). 2mL of 50% slurry beads was used for every 50 mL clarified lysate (i.e., 1 L culture), rotated for 1 hour at 4 °C. Sepharose beads were column captured and separated from the whole lysate by gravity. Beads were washed 2-3 times with lysis buffer. Beads were resuspended in the same initial volume and HRV-3C protease (1 μg per 1 mg expected captured protein) was incubated overnight at 4°C. Cleaved protein was separated from the beads by column-capture. Effective cleavage was checked by PAGE and Coomassie blue stain or stain-free gel. CNOT9 was further purified by gel filtration on a Superdex 200 26/600 column (GE Healthcare) in gel filtration buffer (Tris pH 8 50 mM, NaCl 250 mM, 5% glycerol, TCEP 0.5 mM). Fractions were checked by PAGE and Coomassie blue stain or stain-free gel (BioRad, 4568091). The purified band was also identified by mass spectrometry. Prior to biolayer interferometry (BLI), CNOT9 was desalted with HiPrep^TM^ 26/10 Desalting column and glycerol removed (final buffer: Tris pH 8 50 mM, NaCl 150 mM, TCEP 0.5 mM).

### Biolayer Interferometry

Bio-Layer Interferometry (BLI) experiments were performed in 50 mM Tris pH 8, 150 mM NaCl, 0.5 mM TCEP and 0.05% Tween-20 on an Octet R8 instrument (Fortebio/Sartorius) operating at 25 °C. Octet Streptavidin biosensors were loaded with peptides (0.5 μg/mL) and subsequently exposed to different concentrations of recombinant CNOT9 (6.6-420 μM). Association and dissociation curves were recorded for each concentration.

Data were analysed using the Octet BLI Analysis software (Sartorius) and in-house software (Martin et al., 2021). The equilibrium dissociation constant (Kd) was determined from the instrument response against CNOT9 concentrations using least squares non-linear regression. The equation used was R_eq_=[P_o_]R_max_/[P_o_]+Kd, where R_eq_ is the response at equilibration, P_o_ is protein concentration, and R_max_ is the maximal response when all available binding sites on the sensor are saturated. K_d_ values were calculated as mean values ± standard deviation (n ≥ 3 independent biological replicates).

In control experiments, sensors with no peptides immobilised were exposed to varying CNOT9 concentrations (6.6-420μM), to test for non-specific binding. Response values obtained from control experiments were subtracted from plotted data prior to fitting.

### Alphafold2-multimer and Alphafold3 predictions

In silico structural modelling of protein-protein interactions was performed using Alphafold v2.3.2-multimer, on the local high-performance computing (HPC) cluster at the Francis Crick Institute, utilising one GPU. A total of 25 candidate structural models (five AF2 neural network models, across 5 independent random seeds) were generated and globally ranked by predicted local distance difference test (pLDDT) score and interface predicted template modelling (ipTM). Predicted interactions from a minimum of 5 top ranked predictions, were structurally visualised and analysed using ChimeraX 1.6.1 by their predicted alignment error (PAE) and distance (Å) for consistency and confidence in the prediction. A minimum PAE <15 and distance <5 was used, although more stringent cut-offs were used where possible to increase confidence in predictions. Alphafold2 predictions have been visualised using Pymol^TM^ v3.1.6.1.

Alphafold 3 predictions were generated using the hosted AlphaFold server (alphafoldserver.com). By default, five distinct structural predictions were generated per protein-protein interaction by sampling underlying diffusion process five times from a single random seed. All five structures were ranked by the server’s integrated ranking_score. All five predictions were structurally visualised and analysed using ChimeraX 1.6.1 by their predicted alignment error (PAE) and distance (Å) for consistency and confidence in the prediction. A minimum PAE <15 and distance <5 was used, although more stringent cut-offs were used where possible to increase confidence in predictions. Alphafold3 predictions have been visualised using Pymol^TM^ v3.1.6.1.

In silico screening for candidate interacting proteins with Pho92 (protein of interest) using Alphafold3 was performed using Process_Alphafold3_Outputs (Willich, S. (2024) Process_AlphaFold3_Outputs doi.org/10.5281/zenodo.13925934). PAE<11 and distance (Å) <3 was applied.

### Circular dichromism

Far-UV CD spectra (190-260 nm) were recorded on a Jasco J-815 spectropolarimeter fitted with a cell holder thermostatted by a CDF-426S Peltier unit. Measurements were performed at 20 °C at a peptide concentration of 0.15 mg/ml in 10 mM sodium phosphate pH 8 and 150 mM sodium fluoride, using fused silica cuvettes with 1 mm path length. 2,2,2-Trifluoroethanol (TFE, Sigma-Aldrich) was added at different concentrations (15-50%). Spectra were recorded with 0.2 nm resolution and baseline corrected by subtraction of the appropriate buffer spectrum. CD intensities are presented as the mean residue CD extinction coefficient (Δε_mrw_) calculated as:

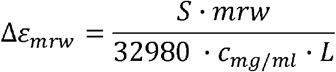

where S is the signal in millidegrees, mrw is the mean residue weight (molecular weight divided by the number of residues), c_mg/ml_ is the concentration in mg/ml, and L is the pathlength (in cm).

### Bioinformatics analysis

FASTQ files were processed using the nf-core/rnaseq pipeline (version 3.10.1; 10.5281/zenodo.1400710), executed using Nextflow (v23.10.0) (Di Tommaso et al., 2017). Reads were aligned to the *Saccharomyces cerevisiae* strain SK1 reference genome (SK1_SGD_2018_NCSL00000000) using STAR (2.7.10a)^52^, and gene-level quantification was performed with RSEM (v1.3.1) ^53^.

Differential expression analysis was carried out with DESeq2 (1.48.1) ^54^ using a simple design formula (∼ group). Wald tests were used to assess differential expression. Multiple-testing correction was performed using Independent Hypothesis Weighting (IHW)^55^; genes with an IHW-adjusted p-value below 0.05 were considered significant. ashr ^56^ was used for effect-size shrinkage.

Functional enrichment analysis was performed using the clusterProfiler (v4.16.0) ^57^ and ReactomePA (v1.52.0) ^58^ packages. Gene Ontology analyses (enrichGO and gseGO) used the *Saccharomyces cerevisiae* organism annotation database org.Sc.sgd.db (3.21.0). Over-Representation Analysis was performed using the enrichGO (Gene Ontology) and enrichPathway (Reactome) functions using genes with FDR < 0.05. Terms with an adjusted p-value ≤ 0.05 after Benjamini–Hochberg correction were considered significant. For Gene Set Enrichment Analysis (GSEA), genes were first ranked using DESeq2’s Wald test statistic and ranked gene lists were then analysed using gseGO and gsePathway functions to identify functionally enriched pathways or gene sets.

Additional GSEA analysis was performed using clusterProfiler’s GSEA function with a custom geneset, ranking genes based on shrunk log-fold changes. GSEA was run for 10,000 permutations and Benjamini–Hochberg correction was applied to identify significantly enriched pathways (adjusted p-value ≤ 0.05).

FASTQ files were processed using the nf-core/rnaseq pipeline (version 3.25.0; 10.5281/zenodo.1400710), executed using Nextflow (v25.10.4) ^59^. Reads were aligned to the *Saccharomyces cerevisiae* strain SK1 reference genome (SK1_SGD_2018_NCSL00000000) using STAR (2.7.11b) ^52^, and gene-level quantification was performed with RSEM (v1.3.1) ^53^.

Differential expression analysis was carried out with DESeq2 (1.50.2) ^54^ using a simple design formula (∼ batch + group). Wald tests were used to assess differential expression. Multiple-testing correction was performed using Independent Hypothesis Weighting (IHW) ^55^; genes with an IHW-adjusted p-value below 0.05 were considered significant. ashr ^56^ was used for effect-size shrinkage. Functional enrichment analysis was performed using the clusterProfiler (v4.18.4) (Wu et al., 2021) and ReactomePA (v1.54.0) ^58^ packages. Gene Ontology analyses (enrichGO and gseGO) used the *Saccharomyces cerevisiae* organism annotation database org.Sc.sgd.db (3.22.0). Over-Representation Analysis was performed using the enrichGO (Gene Ontology) and enrichPathway (Reactome) functions using genes with FDR < 0.05. Terms with an adjusted p-value ≤ 0.05 after Benjamini– Hochberg correction were considered significant. For Gene Set Enrichment Analysis (GSEA), genes were first ranked using DESeq2’s Wald test statistic and ranked gene lists were then analysed using gseGO and gsePathway functions to identify functionally enriched pathways or gene sets.

Additional GSEA analysis was performed using clusterProfiler’s GSEA function with a custom gene set, ranking genes based on shrunk log-fold changes. GSEA was run for 10,000 permutations and Benjamini–Hochberg correction was applied to identify significantly enriched pathways (adjusted p-value ≤ 0.05).

## Statistical analysis

Statistics used for the figures are indicated in the figure legends.

## Data Availability

The accession numbers for the sequencing data reported in this paper are deposited in GEO under GSE339410, GSE339409, and GSE339501.

## Acknowledgements

We acknowledge the Genomics STP, and particularly Deb Jackson, Daniel Leonce, Marg Crawford and Ashley Fowler, for their contributions to mRNA library preparation and sequencing. We acknowledge Avinash Ghanate of the Bioinformatics STP. We acknowledge Raveena Preema, Huda Khalaf and Grant Pellowe for their help in setting up and troubleshooting recombinant protein purification. We thank Dhira Joshi of the Peptide Chemistry STP for the design and production of bespoke peptides. We thank the Flow Cytometry STP for training and access to the FACS instruments. We also thank the members of the van Werven and Kranc lab for the critical reading of the manuscript. We thank Sebastiaan Winkler for sharing his expertise on the CNOT9 protein. We thank Nicolas Ingolia for sharing the dual reporter constructs. We thank Joseph Reese and Martine Collart for sharing yeast Ccr4-NOT antibodies.

---

**Figure S1. m6A-dependent and m6A-independent proximity labelling of Pho92.** (**A**) Western blot detecting expression of Pho92-TID. Shown are wild-type and *PHO92-TID* induced during meiosis (SPO, 4 hours) (FW1511 and FW11105). Membranes were probed with Myc antibodies. Hxk1 was used as a loading control. (**B**) Scheme of purification of biotinylated proteins for proximity labelling (see materials and methods for details). **(C)** Analysis of eluates of proximity labelling with Pho92-TID. Western blots were probed with Myc antibodies to detect Pho92-TID. (**D**) Similar analysis as B except that Western blot membranes were probed with Streptavidin-HRP. (**E**) Onset of meiosis of *PHO92-TID* in wild-type background, and in *pho92*Δ cells as the negative control (FW11105 and FW3528). Cells were induced to enter meiosis. Samples were taken at the indicated time points for DAPI staining. Cells were fixed, stained, and DAPI masses were counted for at least 200 cells per biological repeat. Cells with two or more DAPI masses were considered in meiosis. The mean and SD of n = 3 biological repeats are displayed. (**F**) Gene ontology (GO) analysis of enriched proteins of the proximity labelling experiments of Pho92-TID profiles in the *slz1*Δ background Figure 1C. (**G**) Representative AlphaFold 3 PAE plot of Pho92 and Caf40.

**Figure S2. Pho92 and Caf40/CNOT9 interactions have distinct requirements *in vitro* and in cells.** (**A**) Representative Pho92-Caf40 AlphaFold2 model after review of a minimum of 5 predictions. For Pho92 W5 (cyan), Caf40 (limegreen) V248, Y282 and V293 (magenta) are highlighted as consistent interaction sites across models (PAE<15, Å <5). For Pho92 28-52 (magenta) I28, L32, L35, L38, I39 and L42 are highlighted as consistent interaction sites with Caf40 (limegreen) Y218, L221, L261, T264, V265. (**B**) Representative Pho92-Caf40 Alphafold3 model after review of a minimum of 5 predictions. For Pho92 W5 (cyan), Caf40 (limegreen) L101, Y107, S143 are highlighted as consistent interaction sites across models (PAE<15, Å<5). For Pho92 28-52 (magenta) I28, L35, L38, I39, L42 (orange), Caf40 (limegreen) Y218, L221, L271, V265, T264 (magenta) are highlighted as consistent interaction sites. (**C**) Representative AlphaFold3 model for Pho92 and CNOT9, and its corresponding PAE plot. Highlighted are consistent interaction sites across a minimum of 5 models (PAE<15, Å<5). Pho92 28-52 (magenta), with I28, L32, L35, L38, I39, L42 (orange) highlighted. CNOT9 (limegreen), with G141, K148, T180, V181, F184 (magenta) highlighted. (**D**) Representative Alphafold2 model for Pho92 and CNOT9, and its corresponding PAE plot. Highlighted are consistent interaction sites across a minimum of 5 models (PAE<15, Å<5). Pho92 28-52 (magenta), with I28, L32, L35, L38, I39, L42 (orange). CNOT9 (limegreen), with Y134, L137, G141, K148, L177, V181 (magenta) highlighted. (**E**) Purification of CNOT9 protein. Shown is a Coomassie-stained gel of different fractions of the CNOT9 protein at the final stage of the purification, during size exclusion (see materials and methods for details).

**Figure S3. The IDR and α-helical region are essential for Pho92 effector function. (A)** Western blot of *pCUP-PHO92* and IDR mutants and deletions described in Figure 1C. Pho92 was detected with anti-HA antibodies. As a loading control, Hxk1 was used. (**B**) Reporter signal of λN protein fragment as determined by iRFP for the experiment and constructs described in Figure 3A.

**Figure S4. Caf40 is specifically required for the turnover of m6A-modified transcripts, but not in the tethered reporter.** (**A**) Western blot of the strains and conditions described in A to determine the depletion efficiency of AID strains. Membranes were probed with anti-V5 antibodies and, as a loading control, with anti-Hxk1 antibodies. (**B**) Relative m6A levels determined by m6A-ELISA of strains and conditions described in A. As a control, we included wild-type cells expressing the *TIR1* ligase. (**C**) PCA plot of RNA-seq samples described in Figure 3. (**D**) RNA-seq analysis of RNA regulatory depletion strains. Shown are boxplots of Pho92 targets to which Pho92 associates in an m6A-dependent manner. We also included a control set of transcripts that served as a negative control. The RNA-seq signal of the depletions (AID + IAA) was normalised to the wild-type expressing the TIR ligase (+IAA). The RNA-seq signal for *pho92*Δ was directly compared with the wild-type control. The box plots are sorted by higher or lower mean signal in Pho92 targets relative to the control.

**Figure S5. Pho92 IDR is promiscuous in a tethered reporter context. (A)** AlphaFold 3 PAE plot of Pho92 and Not1**. (B)** Same as (A), but Not1 and Caf40. **(C)** Same as (A), but Pho92 and *not1-*Δ*duf1.* (**D**) Same as (A), but Caf40 and *not1-* Δ*duf1*.

**Figure S6. A single hydrophobic residue is essential for Pho92 effector function. (A)** RNA levels by qPCR. Shown are empty vector control (EV), full-length Pho92 (FL), the N-terminal IDR (N, residues 1 to 155), and N-terminal IDR with L35A and L38P (FW855, FW938, FW939). The mean signal (and SD) relative to the EV control is shown across 3 biological replicates. (**B**) Circular Dichroism (CD) spectroscopy of Pho92_28-52_, Pho92_28-52, L35A_, and Pho92_28-52, 4A_ peptides in the presence of increasing concentrations of 2,2,2-trifluoroethanol (TFE). (**C**) Western blot of *pCUP-PHO92* and *pho92* mutants. Pho92 was detected with anti-HA antibodies. As a loading control, Hxk1 was used.

**Figure S7. YTHDFs’ IDR hydrophobic patches are required for effector activity.** (**A-B**) Structural disorder predictions using IUPrep2 (beige) and pLDDT (AlphaFold2) blue scores for YTHDF1 (A) and YTHDF2 (B). The x-axis indicates the residue position, and the y-axis the score. **(C)** AlphaFold2 PAE plot of YTHDF2 and CNOT9**. (D)** AlphaFold2 PAE plot of YTHDF2 and Caf40. **(E)** AlphaFold2-multimer YTHDF2 and Caf40 interaction. The YTHDF2 W75 predicted interaction is highlighted with consistent predicted Caf40 residues V248, P249, L252, Y282, R289, A292, V293, minimum (PAE <15, Å<5) **(F)** AF3 prediction of YTHDF2 and CNOT9; the YTHDF2 W134 interaction is highlighted. Consistent and conserved CNOT9 residues I164, P165, L168, Y198, R205, H208, V209 are highlighted (PAE <15, Å<5). **(G)** AlphaFold3 prediction of YTHDF2 and Caf40; the YTHDF2 W134 interaction is highlighted. Consistent and conserved Caf40 residues V248, P249, L252, Y282, R289, A292 are highlighted, minimum (PAE <15, Å<5). **(H)** AF3 PAE plot of YTHDF2 and CNOT9. **(I)** AlphaFold3 PAE plot of YTHDF2 and Caf40.

**Figure S8. YTHDFs’ IDR hydrophobic patches are required for effector activity. (A)**Tethered reporter assay of YTHDF2 N-terminal fragment residues 50 to 150 with scanning replacements of 5 residues of linker sequence across the region, totalling 20 constructs. The mean signal (and SD) relative to the empty vector (EV) control is shown across 3 biological replicates. (**B**) Structural disorder prediction using IUPrep2 for the 50-250 region for YTHDF2: WT, LIVFtoST, cLIVFtoST, 4pLIVFtoST, and SCRMBL as described in Figure 7E.

