## Supplementary figures and images for "Cellular context restricts a promiscuous m6A reader IDR to a single functional effector interface for mRNA decay"

### Figures S1-S8

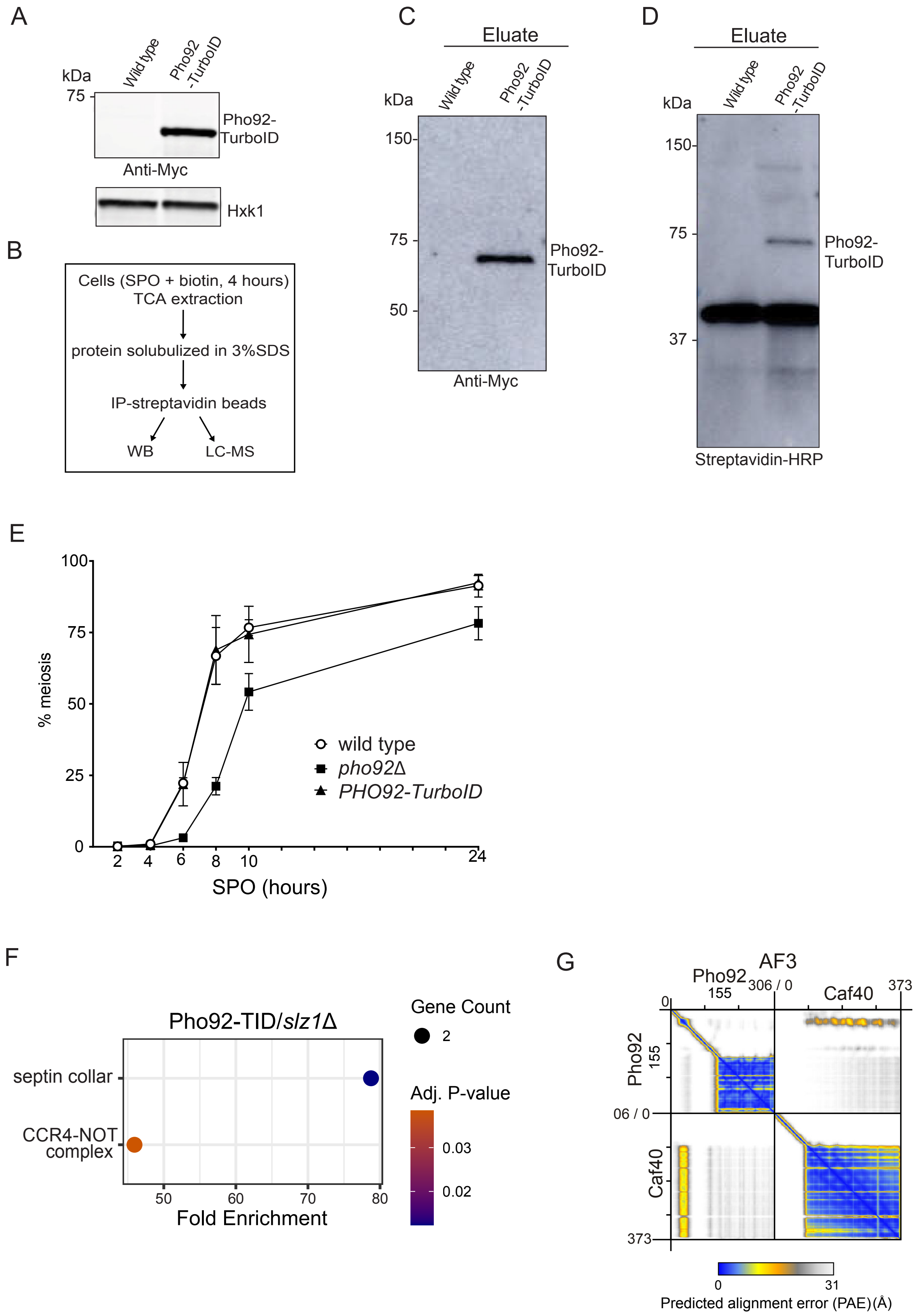

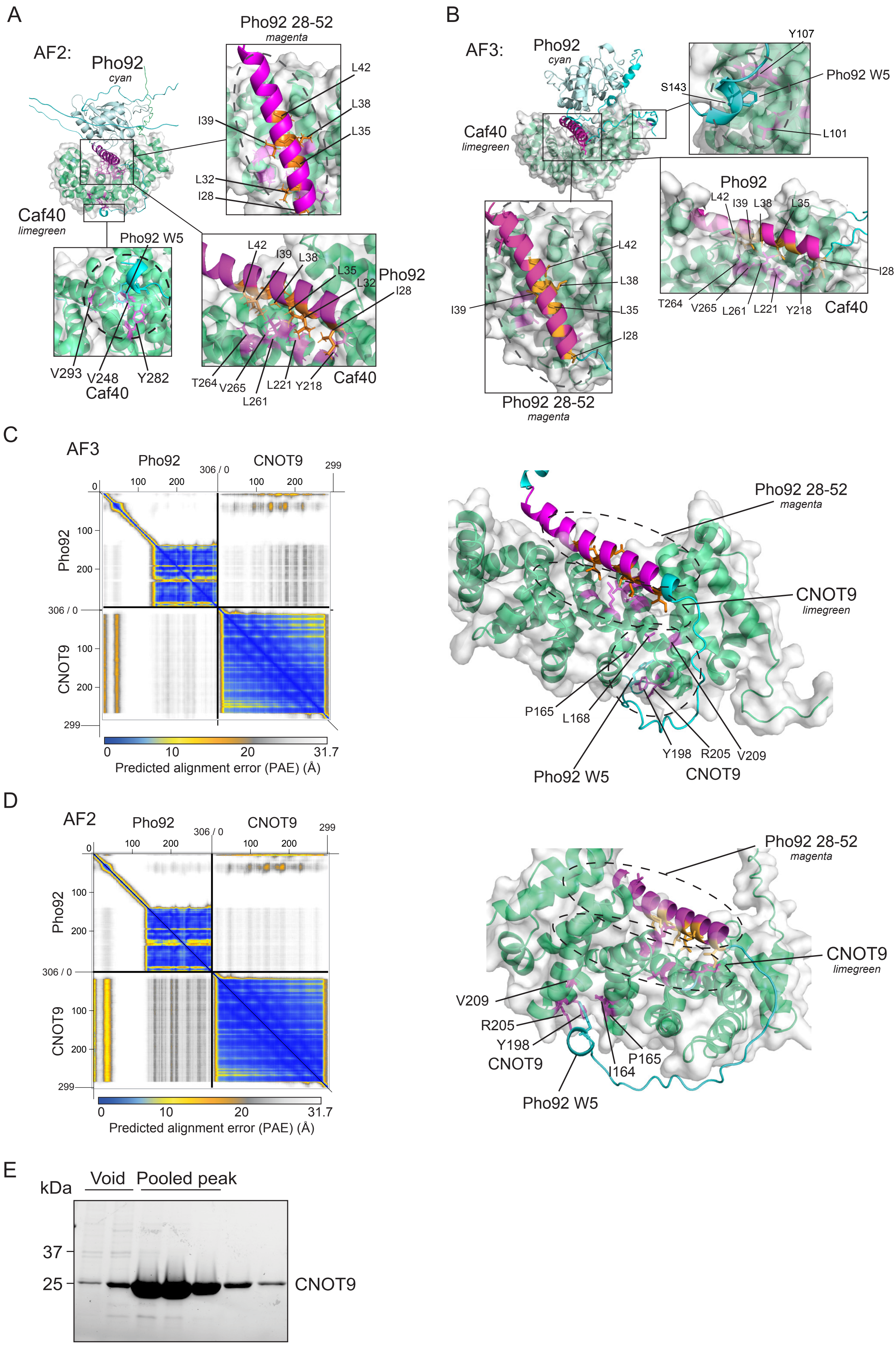

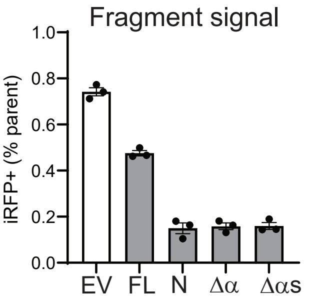

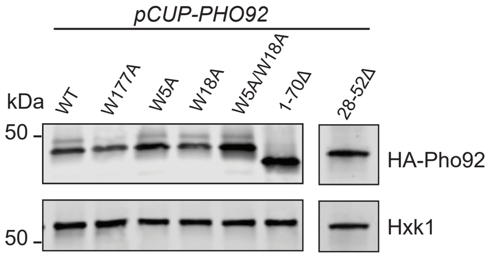

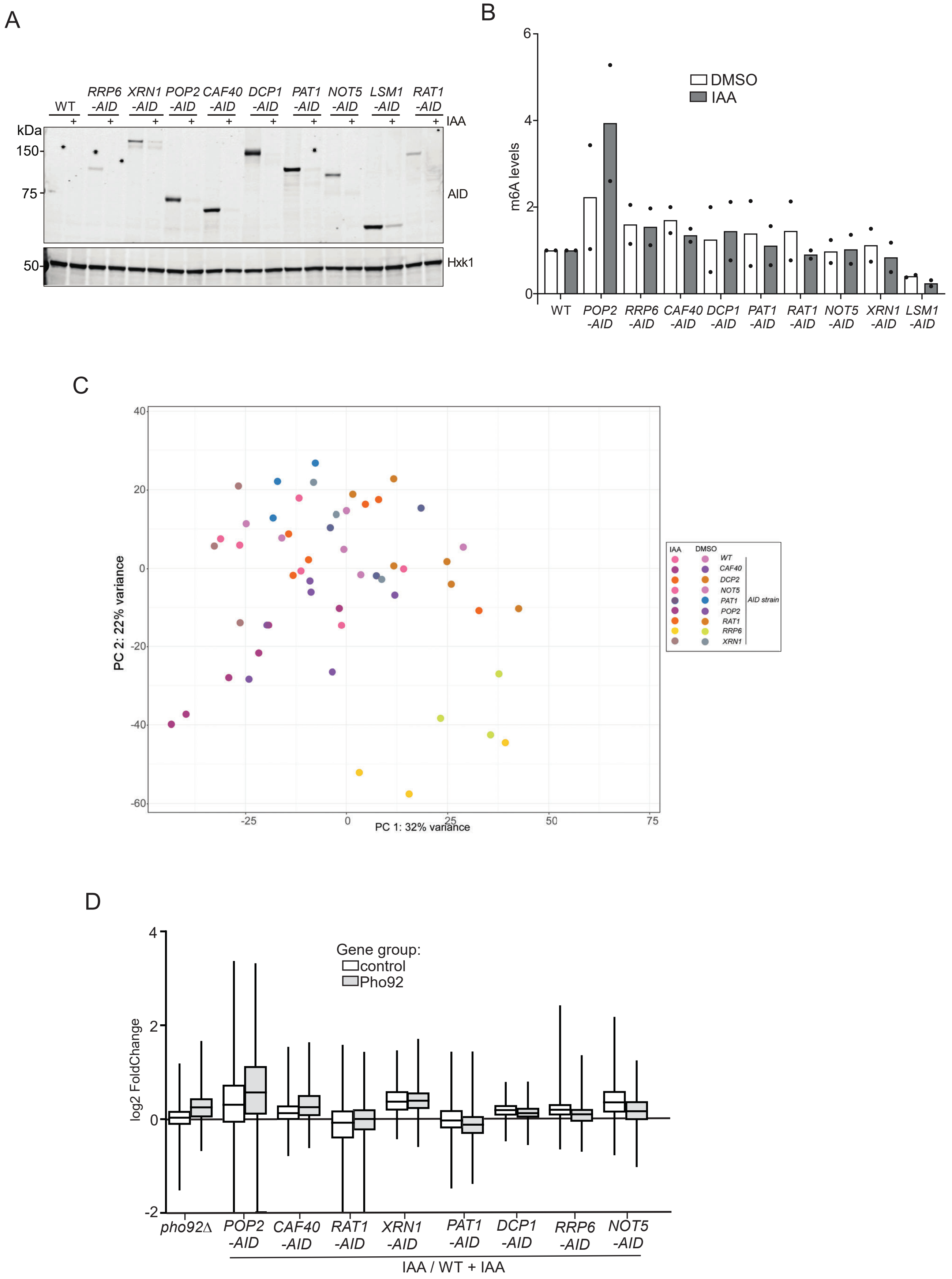

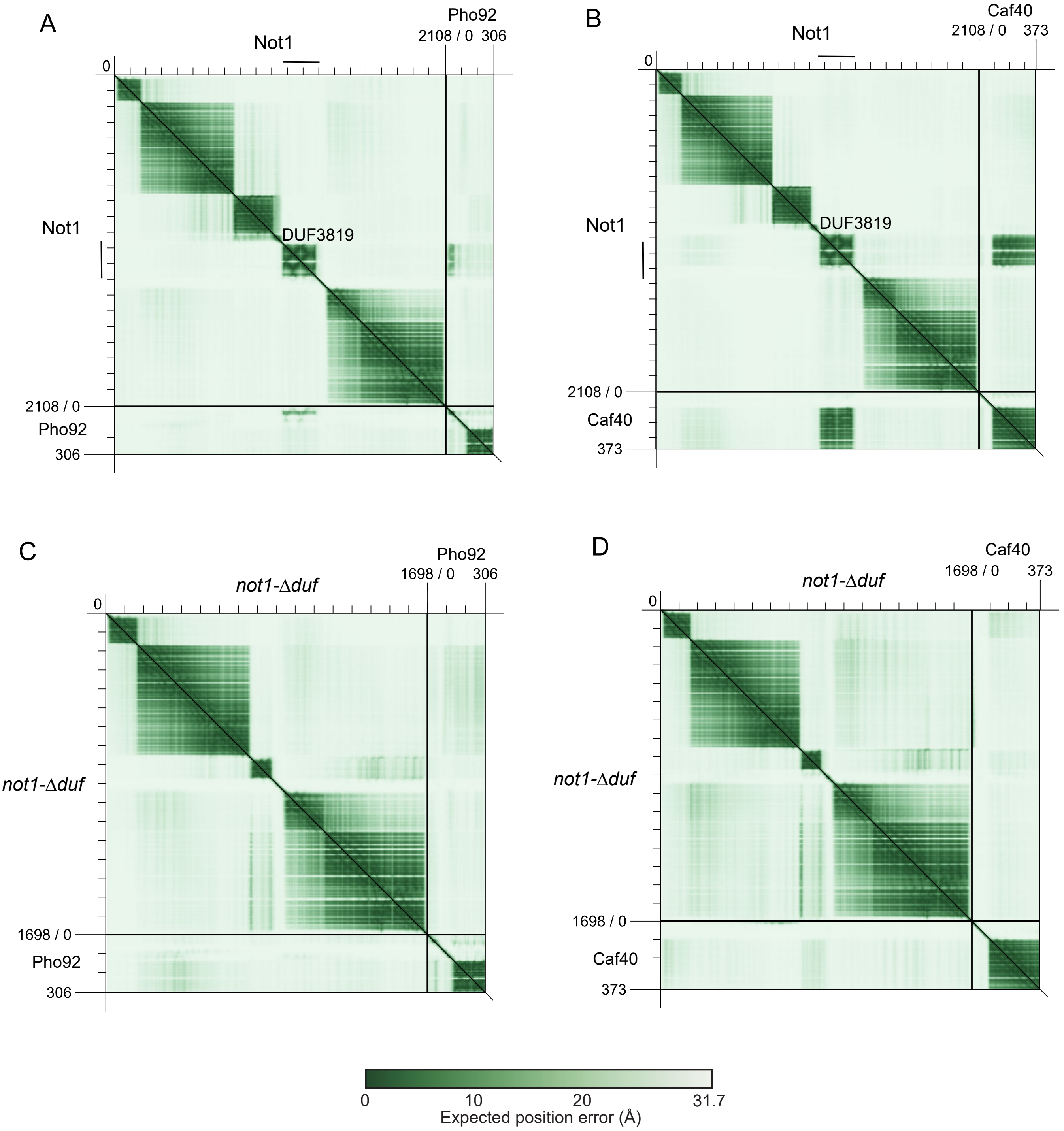

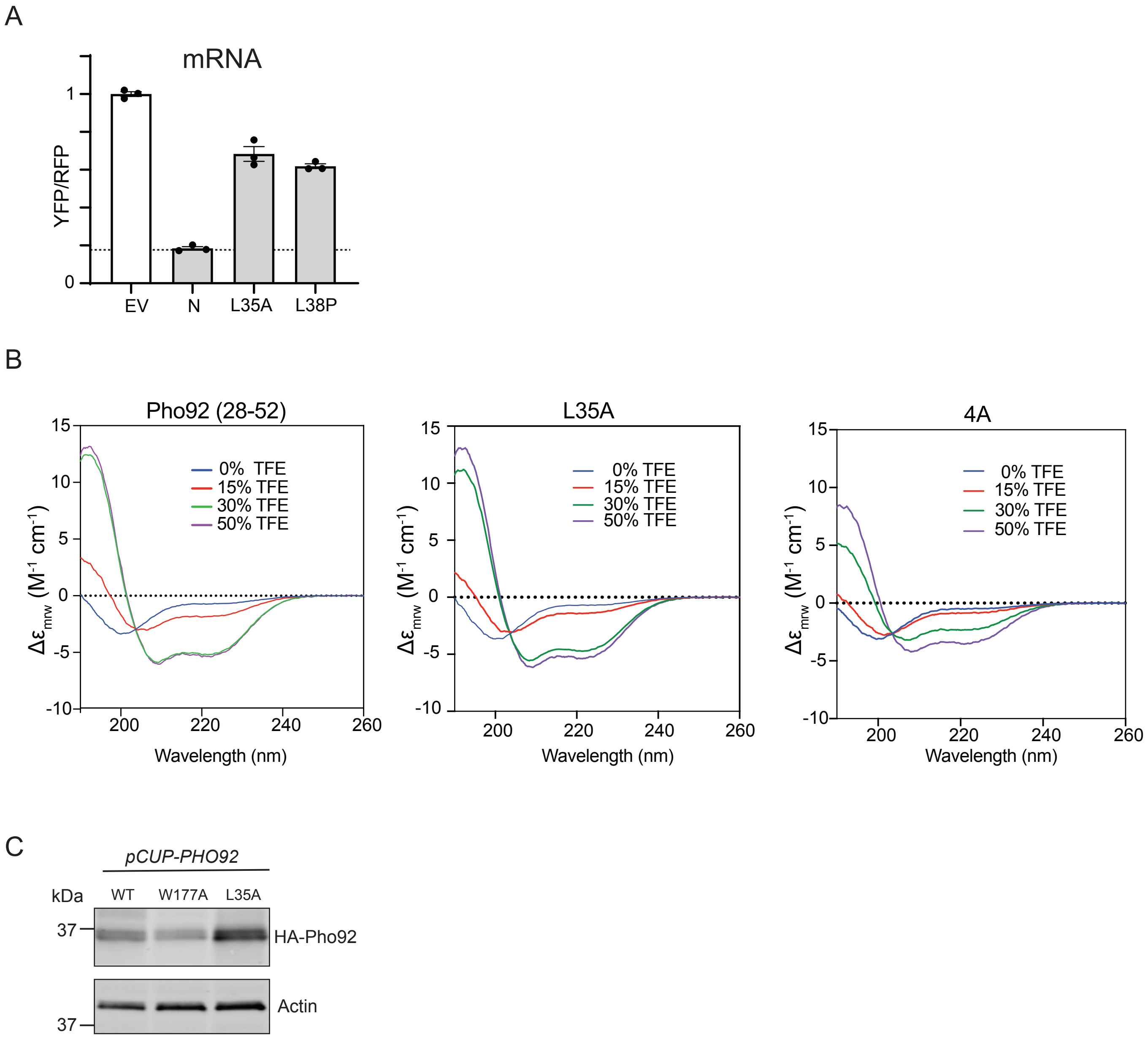

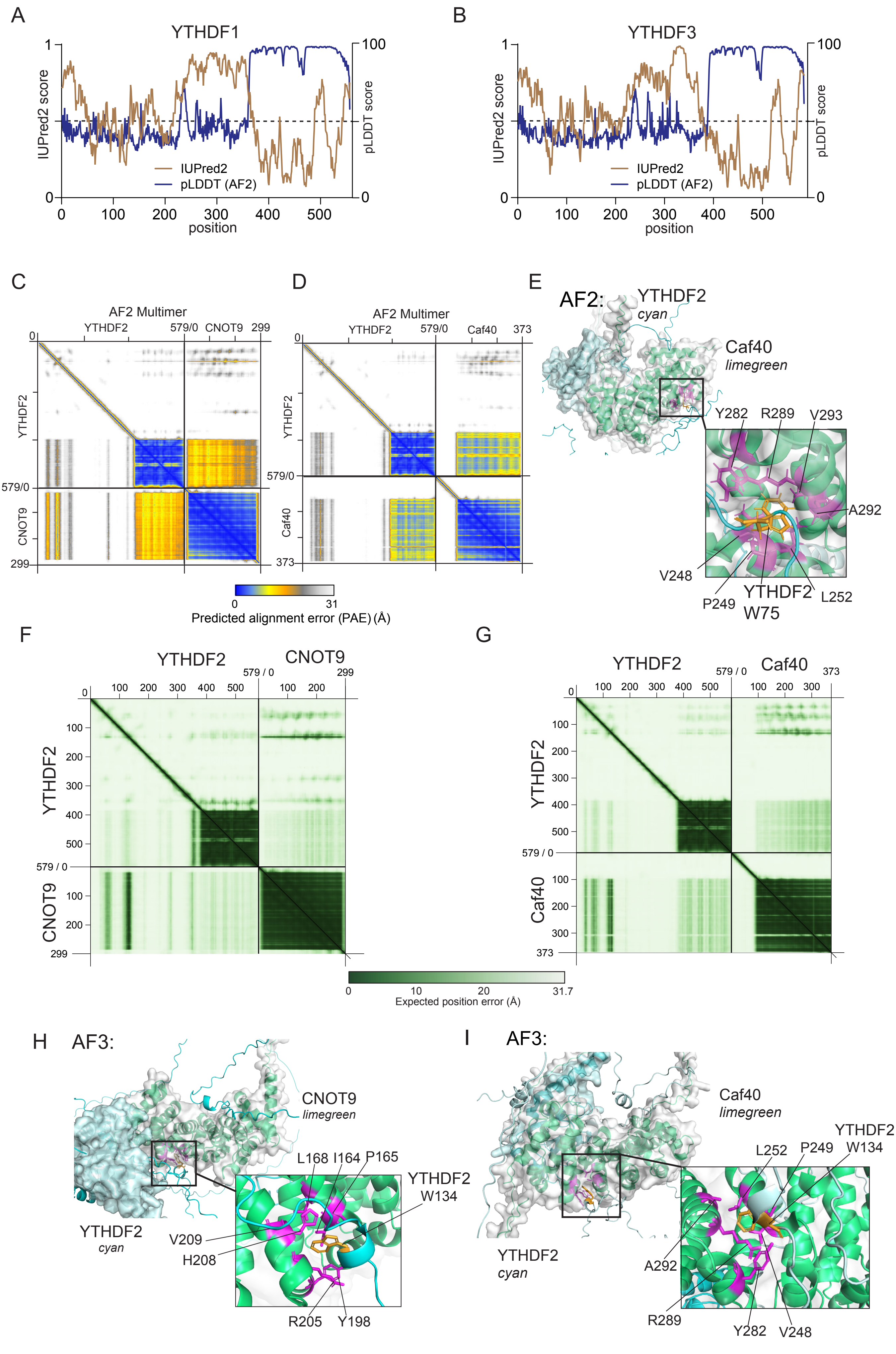

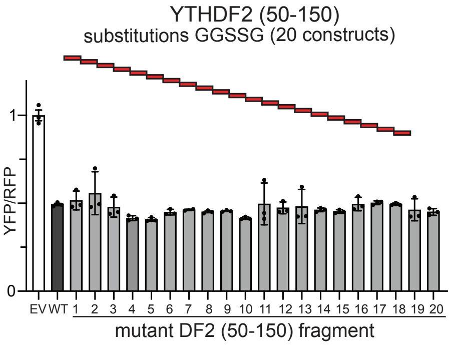

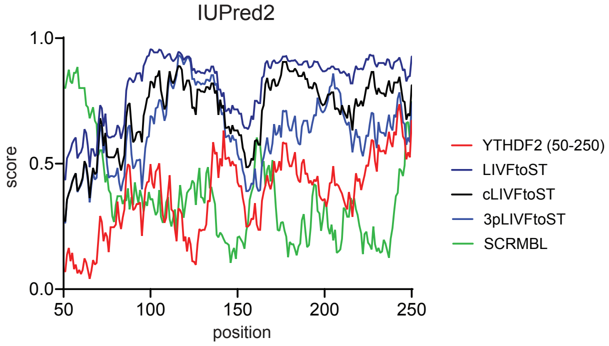
