## Supplementary material for "Cellular context restricts a promiscuous m6A reader IDR to a single functional effector interface for mRNA decay": Tables S1-S4

**Table S1. BLI/Octet, summary equilibrium dissociation constant (Kd) with CNOT9.**

| | | μM | | | | μM | |
|---|---|---|---|---|---|---|---|
| | Peptide | Kd | St dev | | Peptide | Kd | St dev |
| Pho92 | 1-25 | **15** | **2** | YTHDF2 | 124-144 | **107** | **12** |
| Pho92 | 1-25 W5A | **400** | **50** | YTHDF2 | 124-144 W134A | **No binding** | |
| Pho92 | 1-25 W18A | **1220** | **160** | YTHDF2 | 65-85 | **104** | **10** |
| Pho92 | 1-25 W5A/W18A | **> 3000** | | YTHDF2 | 65-85 W75A | **No binding** | |
| Pho92 | 28-52 | **155** | **21** | YTHDF2 | 100-120 | **> 1800** | |
| Pho92 | 28-52 L35A | **1410** | **140** | YTHDF2 | 100-120 L111A/L117A | **No binding** | |
| Pho92 | 28-52_4A | **No binding** | | | | | |

**Table S2. Sequences of YTHDF2 used for reporter experiments.**

**YTHDF2 (residues 50-250)**
MSDSYLPSYYSPSIGFSYSLGEAAWSTGGDTAMPYLTSYGQLSNGEPHFLPDAMF
GQPGALGSTPFLGQHGFNFFPSGIDFSAWGNNSSQGQSTQSSGYSSNYAYAPSSL
GGAMIDGQSAFANETLNKAPGMNTIDQGMAALKLGSTEVASNVPKVVGSAVGSGSI
TSNIVASNSLPPATIAPPKPASWADIASKPAKQQ

**LIVFtoST**
MSDSYTPSYYSPSSGTSYSTGEAAWSTGGDTAMPYSTSYGQSSNGEPHTSPDAM
TGQPGATGSTPTSGQHGSNTSPSGSDTSAWGNNSSQGQSTQSSGYSSNYAYAPS
SSGGAMTDGQSATANETSNKAPGMNTSDQGMAASKSGSTETASNSPKSTGSATG
SGSTTSNSTASNSSPPATSAPPKPASWADSASKPAKQQ

**cLIVFtoST**
MSDSYLPSYYSPSSGTSYSTGEAAWSTGGDTAMPYSTSYGQLSNGEPHFLPDAM
TGQPGATGSTPFSGQHGSNTSPSGIDTSAWGNNSSQGQSTQSSGYSSNYAYAPS
SSGGAMIDGQSATANETSNKAPGMNTSDQGMAASKSGSTEVASNVPKVTGSATGS
GSITSNSVASNSLPPATSAPPKPASWADIASKPAKQQ

**3pLIVFtoST**
MSDSYLPSYYSPSSGSSYSSGEAAWSTGGDTAMPYLTSYGQLSNGEPHFLPDAMF
GQPGATGSTPSTGQHGSNTSPSGSDTSAWGNNSSQGQSTQSSGYSSNYAYAPSS
LGGAMIDGQSAFANETLNKAPGMNTIDQGMAALKLGSTESASNSPKTSGSASGSG
SITSNIVASNSLPPATIAPPKPASWADIASKPAKQQ

**SCRMBL**
SPDALSTPGGSQEAKPGDISGSAQLVQMDFQSAMGLQYGSPGSVPLSLNKSAMDI
VAGAAGKANPIYNGPFATSLEPWIDQNGGGVNAFSNQSISFSYYVNGSSSYTPWTG
KTAGPSELTQSKLNYADASIYYGGTSALGSDAAAPKSMFHGPMSYSINGSFSTWNL
TGGPSAPPAIVFAAFSTMFSPLAQGHPNSLQSAE

**Table S3. Genotypes of strains.**

| Lab reference | Genotype |
|---|---|
| FW1511 | *MATa/alpha ho::LYS2, lys2, ura3, leu2::hisG, his3::hisG, trp1::hisG* |
| FW5730 | *MATa/alpha, ho::LYS2, lys2, ura3, leu2::hisG, trp1::hisG his3:p550 RRP6-3V5-IAA7:KanMX6* |
| FW11725 | *MATa/alpha, ho::LYS2, lys2, ura3, leu2::hisG, his3::hisG, trp1::hisG caf40::HYG* |
| FW3528 | *MATa/alpha, ho::LYS2, lys2, ura3, leu2::hisG, his3::hisG, trp1::hisG pho92::HYG* |
| FW3525 | *MATa/alpha, ho::LYS2, lys2, ura3, leu2::hisG, his3::hisG, trp1::hisG Pho92-3V5::KAN-MX* |
| FW5737 | *MATa/alpha, ho::LYS2, lys2, ura3, leu2::hisG, trp1::hisG his3:p550* |
| FW5745 | *MATa/alpha, ho::LYS2, lys2, ura3, leu2::hisG, trp1::hisG his3:p550 XRN1-3V5-IAA7:KanMX6* |
| FW5953 | *MATa/alpha, ho::LYS2, lys2, ura3, leu2::hisG, trp1::hisG his3:p550 POP2-3V5-IAA7:KanMX6* |
| FW5958 | *MATa/alpha, ho::LYS2, lys2, ura3, leu2::hisG, trp1::hisG his3:p550* |
| FW6045 | *MATa/alpha, ho::LYS2, lys2, ura3, leu2::hisG, trp1::hisG his3:p550 DCP2-3V5-IAA7:KanMX6* |
| FW6062 | *MATa/alpha, ho::LYS2, lys2, ura3, leu2::hisG, trp1::hisG his3:p550 PAT1-3V5-IAA7:KanMX6* |
| FW6067 | *MATa/alpha, ho::LYS2, lys2, ura3, leu2::hisG, trp1::hisG his3:p550 NOT5-3V5-IAA7:KanMX6* |
| FW6072 | *MATa/alpha, ho::LYS2, lys2, ura3, leu2::hisG, trp1::hisG his3:p550* |

| | |
|---|---|
| | *LSM1-3V5-IAA7:KanMX6* |
| FW7144 | *MATa/alpha, ho::LYS2, lys2, ura3, leu2::hisG, trp1::hisG*<br>*his3:p550*<br>*RAT1-3V5-IAA7:KanMX6* |
| FW11105 | *MATa/alpha, ho::LYS2, lys2, ura3, leu2::hisG, his3::hisG*<br>*trp1::hisG*<br>*Nat::Pho92-Turbo-ID* |
| FW11200 | *MATa/alpha, ho::LYS2, lys2, ura3, leu2::hisG, his3::hisG*<br>*trp1::hisG*<br>*Nat::Pho92-TurboID*<br>*Nat::slz1Δ* |
| FW12148 | *MATa/alpha, ho::LYS2, lys2, ura3, leu2::hisG, his3::hisG*<br>*trp1::hisG*<br>*Nat::Pho92-TurboID*<br>*Hyg::caf40Δ* |
| FW10829 | *MATa/alpha, ho::LYS2, lys2, ura3, leu2::hisG, his3::hisG*<br>*trp1::hisG hyg::pho92*<br>*Trp1::CUP1-HA-PHO92-WT* |
| FW11775 | *MATa/alpha, ho::LYS2, lys2, ura3, leu2::hisG, his3::hisG*<br>*trp1::hisG*<br>*Trp1::CUP1-HA-PHO92-WT*<br>*G418::CAF40-V5* |
| FW11951 | *MATa/alpha, ho::LYS2, lys2, ura3, leu2::hisG, his3::hisG*<br>*trp1::hisG*<br>*Trp1::CUP1-HA-PHO92-W177A*<br>*G418::CAF40-V5* |
| FW11938 | *Mat a/alpha ho::LYS2, lys2, ura3, leu2::hisG, his3::hisG,*<br>*trp1::hisG*<br>*hyg::pho92*<br>*trp1::pCUP-HA-Pho92-W18A*<br>*G418::CAF40-V5* |
| FW11945 | *Mat a/alpha ho::LYS2, lys2, ura3, leu2::hisG, his3::hisG,*<br>*trp1::hisG*<br>*hyg::pho92*<br>*trp1::pCUP-HA-Pho92-W5A*<br>*G418::CAF40-V5* |
| FW11971 | *Mat a/alpha ho::LYS2, lys2, ura3, leu2::hisG, his3::hisG,*<br>*trp1::hisG*<br>*hyg::pho92*<br>*trp1::pCUP-HA-Pho92-W5A/W18A*<br>*G418::CAF40-V5* |
| FW12128 | *MATa/alpha, ho::LYS2, lys2, ura3, leu2::hisG, his3::hisG*<br>*trp1::hisG*<br>*hyg::pho92*<br>*trp1::CUP1-HA-PHO92-1-60Δ*<br>*G418::CAF40-V5* |

| | |
|---|---|
| FW12068 | *MATa/alpha, ho::LYS2, lys2, ura3, leu2::hisG, his3::hisG*<br>*trp1::hisG*<br>*hyg::pho92*<br>*trp1::CUP1-HA-PHO92-1-70Δ*<br>*G418::CAF40-V5* |
| FW12254 | *MATa/alpha, ho::LYS2, lys2, ura3, leu2::hisG, his3::hisG*<br>*trp1::hisG*<br>*hyg::pho92*<br>*trp1::CUP1-HA-PHO92-W5A/W18A/28-52Δ*<br>*G418::CAF40-V5* |
| FW12288 | *MATa/alpha, ho::LYS2, lys2, ura3, leu2::hisG, his3::hisG*<br>*trp1::hisG*<br>*hyg::pho92*<br>*trp1::CUP1-HA-PHO92-28-52Δ*<br>*G418::CAF40-V5* |
| FW11725 | *MATa/alpha, ho::LYS2, lys2, ura3, leu2::hisG, his3::hisG*<br>*trp1::hisG*<br>*hyg::caf40* |
| FW10892 | *MATa/alpha, ho::LYS2, lys2, ura3, leu2::hisG, his3::hisG*<br>*trp1::hisG*<br>*hyg::pho92*<br>*trp1::CUP1-HA-PHO92-W177A* |
| FW11824 | *MATa/alpha, ho::LYS2, lys2, ura3, leu2::hisG, his3::hisG*<br>*trp1::hisG*<br>*hyg::pho92*<br>*trp1::CUP1-HA-PHO92-W5A* |
| FW11830 | *MATa/alpha, ho::LYS2, lys2, ura3, leu2::hisG, his3::hisG*<br>*trp1::hisG*<br>*hyg::pho92*<br>*trp1::CUP1-HA-PHO92-W18A* |
| FW11878 | *MATa/alpha, ho::LYS2, lys2, ura3, leu2::hisG, his3::hisG*<br>*trp1::hisG*<br>*hyg::pho92*<br>*trp1::CUP1-HA-PHO92-W5A/W18A* |
| FW12069 | *MATa/alpha, ho::LYS2, lys2, ura3, leu2::hisG, his3::hisG*<br>*trp1::hisG*<br>*hyg::pho92*<br>*trp1::CUP1-HA-PHO92-1-70Δ* |
| FW12439 | *MATa/alpha, ho::LYS2, lys2, ura3, leu2::hisG, his3::hisG*<br>*trp1::hisG*<br>*hyg::pho92*<br>*trp1::CUP1-HA-PHO92-28-52Δ* |
| FW12437 | *MATa/alpha, ho::LYS2, lys2, ura3, leu2::hisG, his3::hisG*<br>*trp1::hisG*<br>*hyg::pho92*<br>*trp1::CUP1-HA-PHO92-W5A/W18A-28-52Δ* |
| FW12912 | *MATa/alpha, ho::LYS2, lys2, ura3, leu2::hisG, his3::hisG*<br>*trp1::hisG*<br>*hyg::pho92*<br>*trp1::CUP1-HA-PHO92-L35A* |

| | |
|---|---|
| FW12964 | *MATa/alpha, ho::LYS2, lys2, ura3, leu2::hisG, his3::hisG*<br>*trp1::hisG*<br>*hyg::pho92*<br>*trp1::CUP1-HA-PHO92-L35-42 Δ* |
| FW11473 | *MATa/alpha, ho::LYS2, lys2, ura3, leu2::hisG, his3::hisG*<br>*trp1::hisG*<br>*hyg::pho92*<br>*trp1::CUP1-HA-YTHDF2* |
| FW11991 | *MATa/alpha, ho::LYS2, lys2, ura3, leu2::hisG, his3::hisG*<br>*trp1::hisG*<br>*hyg::pho92*<br>*trp1::CUP1-HA-YTHDF2-W422A* |
| FW12877 | *MATα his3Δ1 leu2Δ0 met15Δ0 ura3Δ0*<br>*his3::citrine-boxB*<br>*leu2::mscarlet* |
| FW13065 | *MATα his3Δ1 leu2Δ0 met15Δ0 ura3Δ0 duf3819Δ*<br>*his3::citrine-boxB*<br>*leu2::mscarlet* |
| FW13091 | *MATa his3Δ1 leu2Δ0 met15Δ0 ura3Δ0 duf3819Δ*<br>*his3::citrine-boxB*<br>*leu2::mscarlet*<br>*caf40Δ::natMX* |
| FW13113 | *MATa his3Δ1 leu2Δ0 met15Δ0 ura3Δ0*<br>*his3::citrine-boxB*<br>*leu2::mscarlet*<br>*caf40Δ::natMX* |
| HC053 | *MATα his3Δ1 leu2Δ0 met15Δ0 ura3Δ0*<br>*his3::citrine-boxB*<br>*leu2::mscarlet*<br>*ccr4Δ::natMX* |
| HC060 | *MATα his3Δ1 leu2Δ0 met15Δ0 ura3Δ0*<br>*his3::citrine-boxB*<br>*leu2::mscarlet*<br>*pop2Δ::natMX* |
| EJ401 | *MATα his3Δ1 leu2Δ0 met15Δ0 ura3Δ0 duf3819Δ*<br>*his3::citrine-boxB*<br>*leu2::mscarlet*<br>*ura3::emi-RFP670-lambda-N(empty)*<br>*caf40Δ::natMX* |

**Table S4. Plasmid used.**

| **Lab Reference** | **Description** |
|---|---|
| FW845 | P(PGK1)-yeCitrine-5xBoxB-t(ADH1) SpHis5 |
| FW846 | P(PGK1)-mScarlet-5xPP7-t(CYC1) KlLeu2 |

| | |
|---|---|
| FW847 | P(PGK1)-emiRFP670 t(cyc1) CEN/ARS - empty |
| FW848 | P(PGK1)-emiRFP670 t(cyc1) CEN/ARS – Pho92-17-66 |
| FW855 | P(PGK1)-emiRFP670 t(cyc1) CEN/ARS – Pho92-1-155 (N) |
| FW856 | P(PGK1)-emiRFP670 t(cyc1) CEN/ARS – N W5A |
| FW857 | P(PGK1)-emiRFP670 t(cyc1) CEN/ARS – N W18A |
| FW858 | P(PGK1)-emiRFP670 t(cyc1) CEN/ARS – N W5A/W18A |
| FW859 | P(PGK1)-emiRFP670 t(cyc1) CEN/ARS – N 28-52Δ |
| FW916 | P(PGK1)-emiRFP670 t(cyc1) CEN/ARS – Pho92-1-306 (FL) |
| FW937 | P(PGK1)-emiRFP670 t(cyc1) CEN/ARS – N 35-42Δ |
| FW939 | P(PGK1)-emiRFP670 t(cyc1) CEN/ARS – N L35A |
| FW876 | P(PGK1)-emiRFP670 t(cyc1) CEN/ARS – N L38A |
| FW938 | P(PGK1)-emiRFP670 t(cyc1) CEN/ARS – N L38P |
| FW877 | P(PGK1)-emiRFP670 t(cyc1) CEN/ARS – N I39A |
| FW940 | P(PGK1)-emiRFP670 t(cyc1) CEN/ARS – N L42A |
| FW941 | P(PGK1)-emiRFP670 t(cyc1) CEN/ARS – N L35A/L38A/I39A/L42A (4A) |
| FW967 | P(PGK1)-emiRFP670 t(cyc1) CEN/ARS – Pho92-28-52 |
| FW969 | P(PGK1)-emiRFP670 t(cyc1) CEN/ARS – Pho92-35-42 |
| FW1068 | P(PGK1)-emiRFP670 t(cyc1) CEN/ARS – YTHDF1 1-388 (YTHDF1 N) |
| FW1070 | P(PGK1)-emiRFP670 t(cyc1) CEN/ARS – YTHDF3 1-415 (YTHDF3 N) |
| FW897 | P(PGK1)-emiRFP670 t(cyc1) CEN/ARS – YTHDF2 1-410 (YTHDF2 N) |

| | |
|---|---|
| FW919 | P(PGK1)-emiRFP670 t(cyc1) CEN/ARS – YTHDF2 1-579 (YTHDF2 FL) |
| FW903 | P(PGK1)-emiRFP670 t(cyc1) CEN/ARS – YTHDF2 100-250 |
| FW904 | P(PGK1)-emiRFP670 t(cyc1) CEN/ARS – YTHDF2 150-250 |
| FW905 | P(PGK1)-emiRFP670 t(cyc1) CEN/ARS – YTHDF2 200-250 |
| FW902 | P(PGK1)-emiRFP670 t(cyc1) CEN/ARS – YTHDF2 50-250 |
| FW1085 | P(PGK1)-emiRFP670 t(cyc1) CEN/ARS – YTHDF2 50-250 (LIVFtoST) |
| FW1086 | P(PGK1)-emiRFP670 t(cyc1) CEN/ARS – YTHDF2 50-250 (cLIVFtoST) |
| FW1088 | P(PGK1)-emiRFP670 t(cyc1) CEN/ARS – YTHDF2 50-250 (3pLIVFtoST) |
| FW1090 | P(PGK1)-emiRFP670 t(cyc1) CEN/ARS – (SCRMBL) |
